# The Balance between Saturated and Unsaturated Fatty Acids Regulates Ovarian Cancer Cell Fate

**DOI:** 10.1101/2022.05.24.493247

**Authors:** Guangyuan Zhao, Yuying Tan, Horacio Cardenas, David Vayngart, Hao Huang, Yinu Wang, Russell Keathley, Jian-Jun Wei, Christina R. Ferreira, Ji-Xin Cheng, Daniela Matei

**Affiliations:** Department of Obstetrics and Gynecology, Feinberg School of Medicine, Northwestern University, Chicago, IL 60611, USA; Driskill Graduate Program in Life Sciences, Feinberg School of Medicine, Northwestern University, Chicago, IL 60611, USA; Department of Biomedical Engineering, Boston University, Boston, MA 02215, USA; Feinberg School of Medicine, Northwestern University, Chicago, IL 60611, USA; Department of Pathology, Feinberg School of Medicine, Northwestern University, Chicago, IL 60611, USA; Robert H. Lurie Comprehensive Cancer Center, Chicago, IL 60611, USA; Bindley Bioscience Center, Purdue University, West Lafayette, IN 47906, USA; Department of Electrical and Computer Engineering, Boston University, Boston, MA 02215; Photonics Center, Boston University, Boston, MA 02215, USA; Jesse Brown VA Medical Center, Chicago, IL 60612, USA

**Keywords:** Ovarian cancer, lipid metabolism, fatty acids, ER stress, SRS imaging

## Abstract

Fatty acids are an important source of energy and a key component of phospholipids in membranes and organelles. Saturated (SFAs) are converted into unsaturated fatty acids (UFAs) by stearoyl Co-A desaturase (SCD), an enzyme highly active in cancer. Here we studied how the balance between SFAs and UFAs maintained by SCD impacts cancer cell survival and tumor progression. SCD depletion or inhibition caused lower levels of UFAs vs. SFAs and altered fatty acyl chain plasticity, as demonstrated by lipidomics and stimulated Raman spectroscopy (SRS). Further, the loss of equilibrium between UFAs and SFAs resulting from SCD knock down triggered endoplasmic reticulum (ER) stress response with brisk activation of IRE1α/XBP1 and PERK/eIF2α/ATF4 axes. Stiff and disorganized ER membrane was visualized by electron microscopy and SRS imaging in cells in which SCD was knocked down. The induction of long-term mild ER stress or short-time severe ER stress by the increased levels of SFAs and loss of UFAs led to cell death. However, ER stress and apoptosis could be readily rescued by supplementation with UFAs and re-equilibration of SFA/UFA levels. The effects of SCD knockdown or inhibition observed *in vitro*, translated into suppression of intraperitoneal tumor growth in xenograft models. Furthermore, a combined intervention using an SCD inhibitor and an SFA enriched diet, initiated ER stress in tumors growing *in vivo* and potently blocked their dissemination. In all, our data support SCD as a key regulator of the cancer cell fate under metabolic stress and point to new treatment strategies targeting the lipid balance.

**Significance Statement:** We show that the balance between saturated and unsaturated fatty acids tightly regulated by the desaturase SCD impacts the survival of cancer cells; increased levels of unsaturation being protective against ER stress induced apoptosis. Decreasing fatty acid unsaturation, either through SCD depletion or through SCD inhibition coupled with a dietary intervention blocks tumor progression *in vivo*. Our findings support the concept of targeting the lipid balance as a new target in cancer.

## Introduction

To keep up with the demands of limitless proliferation and metastatic spread, cancer cells thwart physiological metabolic pathways to meet their augmented energetic needs. When glucose and oxygen are in short supply, fat becomes convenient fuel. Whether taken up from the tumor microenvironment (TME) or newly synthesized, lipids function as alternative energy source for rapidly growing tumors. In addition, lipids are key constituents of membranes and intra-cellular organelles, enabling the smooth functioning of signaling circuitries and the homeostasis of cells proliferating under stressful conditions (1). A unique property of ovarian cancer (OC) is its tropism to the omentum, a fat-laden organ, which has been thought to function as feeding soil for rapidly expanding tumors (2, 3). While lipid uptake by cancer cells has been studied to some extent (2-4), the role of *de novo* lipogenesis and of the ensuing balance between unsaturated and saturated lipids remain not fully understood.

Within the *de novo* lipogenesis pathway, lipid desaturation is a key step required for the generation of unsaturated lipids to maintain the membrane fluidity, the integrity of cellular signaling, and the lipid pool for β-oxidation (5). Fatty acid (FA) desaturases catalyze the addition of double carbon bonds in acyl chains, regulating the formation of mono- and poly-unsaturated FAs (MUFA and PUFA, respectively). Stearoyl-CoA desaturase (SCD) converts saturated (SFA) to mono-unsaturated FAs, palmitic and stearic acid to palmitoleic and oleic acid, respectively. SCD is upregulated in cancer (6) and its inhibition was shown to suppress cancer cell proliferation, in conditions depleted of exogenous lipids (7). SCD was also shown to protect cancer cells from lipid peroxidation leading to ferroptosis, through a mechanism dependent on the anti-oxidant coenzyme CoQ10 (8). Using stimulated Raman spectroscopy (SRS) (9, 10), which enables analysis of lipid species in rare cell populations, we recently demonstrated that ovarian cancer stem cells (CSCs) are enriched in UFAs (11). We showed that SCD small molecule inhibitors or shRNA-mediated knock down eliminated ovarian CSCs, delaying tumor initiation (11). However, the mechanisms by which SCD regulates cancer cells’ and CSCs’ survival under stressful conditions are not elucidated.

Through a combined lipidomic, transcriptomic and single cell imaging approach, we zoom in on how the balance between UFAs and SFAs regulates cell survival and tumor progression in OC models. We observed that an increase in SFAs caused by addition of SFA to the media, or by SCD depletion or inhibition, led to significant endoplasmic reticulum (ER) stress inducing apoptosis of cancer cells. This ER stress is likely caused by direct effects of SFAs on the ER membrane fluidity, causing activation of the sensor proteins IRE1α and Protein Kinase R-like ER Kinase (PERK) and could be rescued by addition of UFAs and restoration of the required equilibrium. SCD knock-down inhibited tumor progression *in vivo* and pharmacological inhibition of the enzyme coupled with a diet enriched in SFAs had potent anti-tumorigenic effects. These results point to the role of SCD as a gate keeper ensuring the survival of ovarian cancer cells under the continuous metabolic stress imposed by non-stop proliferation and to the therapeutic potential of targeting the lipid balance in cancer.

## Results

### SCD is highly expressed in OC cell lines and tumors

SCD expression was assessed by using representative cancer cell lines, an ovarian cancer TMA, and RNA sequencing data from human specimens profiled by The Cancer Genome Atlas (TCGA) (4) and the Genotype-Tissue Expression (GTEx) Project. *SCD* expression was significantly higher in high grade serous ovarian tumors (HGSOC) profiled by the TCGA (n= 427) compared to normal fallopian tube epithelium (FTE, n = 5, Fig. 1A). Further, *SCD* was highly expressed in OC cell lines as compared to immortalized fallopian tube epithelial cells at both *mRNA* and protein levels (Figs. 1B, C). IHC analysis demonstrated significant upregulation of SCD in HGSOC tumors (n = 12) vs. FTE (n = 6), with 7 of 12 tumors displaying intense staining vs. FTE (Fig. 1D, p = 0.0377). Other histological subtypes of OC also displayed increased SCD expression compared to FTE (p = 0.006), while non-malignant ovarian tumors (borderline tumors and cystadenoma) did not overexpress SCD (Supplementary Table S2).

**Figure 1.**
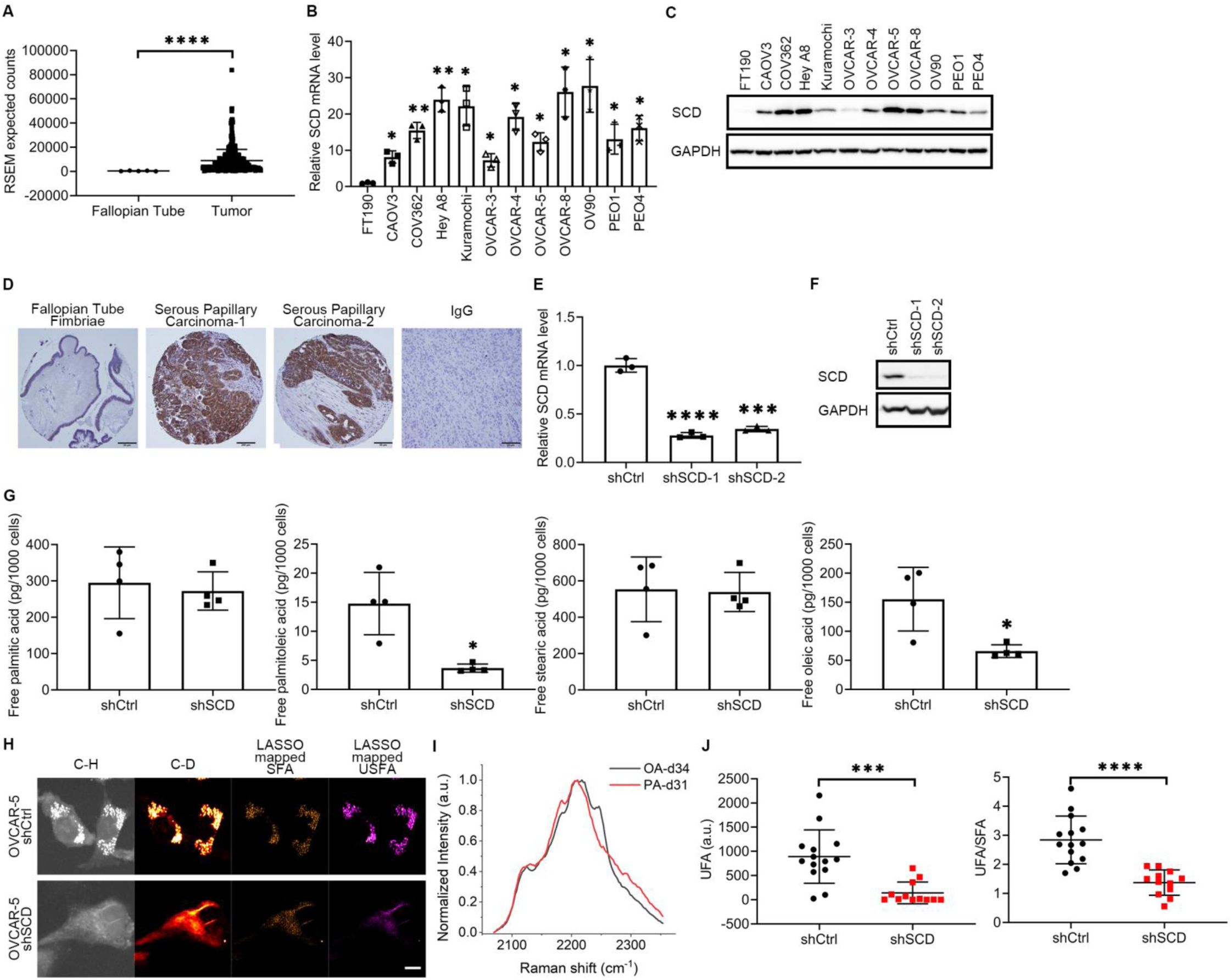
SCD is highly expressed in ovarian cancer and is associated with increased levels of free unsaturated fatty acids. (A) *SCD* expression from RNA-seq data in fallopian tube compared to ovarian cancer tumors in patients from the UCSC Xena Browser. Expression is shown as RSEM expected counts. (B, C) Real-time qRT-PCR analysis of SCD expression (mean ± SD, n = 3) (B), and western blot measurements of SCD protein levels (C) in immortalized fallopian tube luminal epithelial cells (FT190) and eleven ovarian cancer cell lines. (D) Representative images of IHC staining for SCD under 20X magnification in fallopian tube fimbriae (left panel), HGSOC specimens (middle panels). Negative control (IgG) is shown in right panel (tumor tissue). Scale bar corresponds to 50μm. (E) SCD expression measured by qRT-PCR (mean ± SD, n = 3) in OVCAR-5 cells transduced with shRNAs (1 or 2) targeting SCD (shSCD), or control shRNA (shCtrl). (F) Western blot of SCD in shCtrl and shSCD OVCAR-5 cells. (G) Lipidomics analysis of palmitic acid (16:0), palmitoleic acid (16:1), stearic acid (18:0), and oleic acid (18:1) in shCtrl and shSCD OVCAR-5 cells cultured in medium containing low serum (1% FBS) for 48 hours (means ± SD, n = 4). (H) Representative SRS images in C-H and C-D regions, and LASSO mappings of saturated fatty acid (SFA) and unsaturated fatty acids (USFA) in shSCD vs shCtrl OVCAR-5 cells treated with 12.5μM PA-d31 and cultured in low serum conditions. Scale bar: 10 µm. (I) Normalized SRS spectra of oleic acid-d34 (OA-d34) and palmitic acid-d31 (PA-d31) as reference spectrum of UFA and SFA respectively for LASSO analysis. (J) Quantitative analysis of LASSO mapped USFA and ratio between USFA and SFA in shSCD vs shCtrl OVCAR-5 cells. Each data point represents quantitative result from a single cell (n = 12-14). * p < 0.05, ** p < 0.01, *** p < 0.001, **** p < 0.0001.

### SCD regulates the balance between SFAs and UFAs in OC cell

For functional studies, two shRNA sequences targeting *SCD* (shSCD-1 and shSCD-2) or control shRNA (shCtrl) were stably transduced in OVCAR-5, OVCAR-8, OVCAR-3 and PEO1 cells. *SCD* knockdown (KD) was verified by quantitative PCR and western blotting (Figs. 1E, F, Supplementary Figs. S1A-C). Next, lipidomics and isotopic hyperspectral stimulated Raman scattering (hSRS) assessed the abundance of SFAs and UFAs in OC cells transduced with shRNA targeting SCD vs. control shRNA and cultured under low serum conditions to limit the impact of exogenous lipid uptake. A significant reduction in UFAs (palmitoleic and oleic acids), but no difference in abundance of SFAs (palmitic and stearic acids, Fig. 1G) were recorded in cells transduced with shRNA targeting SCD vs. control shRNA. Further, isotopic hSRS imaging compared the levels of UFAs converted from newly imported SFAs in shSCD vs shCtrl transduced cells (Figs. 1H-J). After being maintained in low serum medium for 24 hours, cells were cultured with deuterated SFA (PA-d31) for 24 hours. The C-D bond signal, corresponding to UFAs was measured by hSRS imaging (Fig. 1I), and intracellular deuterated SFA and UFA levels were distinguished through a LASSO unmixing analysis using the distinctive Raman spectra of PA-d31 and OA-d34, as reference for SFA and UFA respectively (Fig. 1H). Images and quantitative analyses of UFA levels and UFA/SFA ratio (Fig. 1J) show that cells in which SCD was knocked down contain significantly less deuterated UFA synthesized from newly imported PA-d31, indicating that the efficiency of converting SFA to UFA is diminished in cells depleted of SCD compared to controls.

To understand which lipid species were most affected by the decrease of UFAs caused by SCD depletion, lipidomics by multiple reaction (MRM) profiling (12) measured the abundance of phosphatidylcholines (PC), phosphatidylethanolamine (PE), sphingomyelin (SM), and triglycerides (TAGs) in OVCAR-5 cells stably transduced with control or shRNA targeting SCD and cultured under low serum conditions for 48 hours. After data normalization, principal component analysis (PCA) confirmed the separation of the two groups (i.e. shCtrl and shSCD) and heatmaps for each lipid species generated by using unsupervised hierarchical clustering demonstrated separation and clustering of the two groups (Supplementary Figure S2). The most affected (fold-change) lipid species by SCD depletion were PC and SM, followed by triacylglycerol (TAG) and PE (Fig. 2A, Supplementary Table S3). A deeper examination of each lipid class revealed that both PC and SM phospholipids with 1° plasticity (one or two carbon-carbon double bonds within the fatty acid tail) (13) were significantly less abundant in shSCD compared with shCtrl transduced cells (p < 0.0001, Fig. 2B). Further, a significant reduction of 16:1 or 18:1 fatty acyl chains integrated TAGs was observed in OC cells stably transduced with shRNA targeting SCD compared to cells transduced with control shRNA (p= 0.0382, Fig. 2C). Combined, the lipidomics analyses coupled with hSRS imaging demonstrate that cells depleted of this key desaturase, contain less abundant UFAs, and less plastic lipid species, particularly PC and SM.

**Figure 2.**
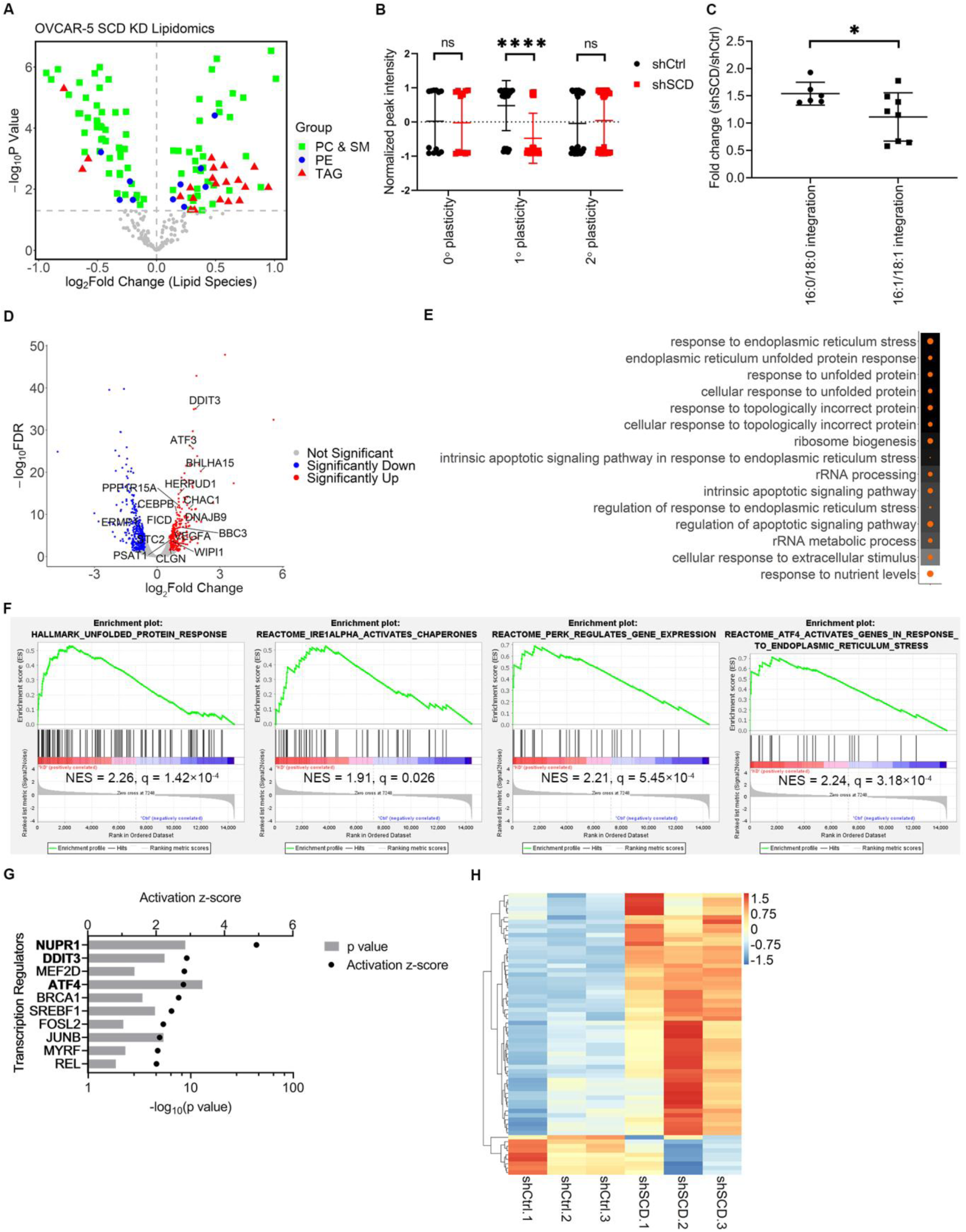
Restricting monounsaturated fatty acids activates the endoplasmic reticulum stress response pathway. (A) Volcano plot of changes in phosphatidylcholine and sphingomyelin (PC & SM, green), phosphatidylethanolamine (PE, blue) and triacylglycerol (TAG, red) relative peak intensities measured by lipidomics profiling analysis in shSCD vs shCtrl OVCAR-5 cells cultured in low serum conditions for 48 hours. (B) Normalized peak intensities of phosphatidylcholine (PC) lipids with differing fatty chain plasticity (0°, no carbon-carbon double bond; 1°, 1 or 2 carbon-carbon double bonds; 2°, more than 2 carbon-carbon double bonds) in shSCD vs shCtrl OVCAR-5 cells. ShSCD cells have significantly fewer 1° PC lipids than shCtrl cells whereas no difference was observed for 0° or 2° PC lipids. (C) Fold change of triacylglycerol (TAG) lipids containing 16:0 or 18:0 fatty acyl chains integration and 16:1 or 18:1 fatty acyl chains integration determined by lipidomics profiling analysis in shSCD vs shCtrl OVCAR-5 cells. (D) Volcano plot of changes in gene expression measured by RNA-seq in shSCD vs shCtrl OVCAR-5 cells. Red dots represent upregulated genes and blue dots, downregulated genes. Genes of the ER stress response pathway are indicated by lines. (E) Dot plot of the top 15 enriched Gene Ontology terms identified by analysis of upregulated genes (RNA-seq) in shSCD vs shCtrl OVCAR-5 cells. Dot size corresponds to ratio of genes on the pathway vs total number of upregulated genes. Background color corresponds to FDR q value. (F) Enrichment plots generated by Gene Set Enrichment Analysis of gene expression (RNA-seq normalized counts) in OVCAR-5 shSCD vs shCtrl using Hallmark and C2 gene sets from Molecular Signatures Database. (G) Top ten significant upstream regulators identified by Ingenuity Pathway Analysis using differentially expressed genes from RNA-seq analysis of OVCAR-5 shSCD vs shCtrl cells. Transcription factors involved in the ER stress response pathway are in bold characters. (H) Heatmap of expression (RNA-seq normalized counts) and supervised hierarchical clustering of significant genes of the ER stress response pathway in shCtrl and shSCD OVCAR-5 cells (n = 3). * p < 0.05, ** p < 0.01, *** p < 0.001, **** p < 0.0001.

### *SCD* knock down triggers ER Stress response

To further understand the global effects of UFA restriction in OC cells, RNA-seq compared the transcriptomic profiles of cells transduced with shRNA targeting SCD or control shRNA. Quality control of RNA-sequencing is included in Supplementary Fig. S3. Cells were cultured under low serum conditions for 48 hours. There were 1513 differentially expressed genes (DEGs; FDR < 0.05), of which 707 were upregulated and 806 downregulated between OVCAR-5 cells stably transduced with control or shRNA targeting SCD (Fig. 2D, Supplementary Table S4). Gene Ontology analysis of significantly upregulated genes revealed that the ER stress response was the top affected pathway by SCD KD (Fig. 2E). Gene Set Enrichment Analysis (GSEA) of all genes also indicated that ER stress response gene sets (Hallmark Unfolded Protein Response; Reactome_IRE1α Activities Chaperones, Reactome PERK Regulates Gene Expression, Reactome ATF4 Activities Genes in Response to Endoplasmic Reticulum Stress) were upregulated in cancer cells in which SCD was knocked down (Fig. 2F). Several genes (*DDIT3, ATF3, PPP1R15A, CEBPB*) involved in ER stress response are highlighted in the volcano plot depicting DEGs between SCD KD and control cells (Fig. 2D). Furthermore, upstream regulator analysis performed with Ingenuity Pathway Analysis (IPA) software identified several transcription factors (*ATF4, DDIT3, NUPR1*) involved in the ER stress response pathway among the top ones predicted to be activated in OC cells transduced with shRNA targeting *SCD* vs control cells (Fig. 2G). A heatmap including significant genes in the ER stress response pathway (n = 64 genes) illustrates clear differences in expression levels for these transcripts between cells in which SCD was knocked down vs. control cells (Fig. 2H). Further, exploration of the transcriptomic dataset associated with the Cancer Dependence Map (14) and Cancer Cell Line Encyclopedia (15) shows that dependency score of key transcription regulators in the ER stress response pathway such as *ATF4* and *DDIT3* were positively correlated with that of SCD (r=0.3117 and 0.3878, respectively Supplementary Figs. S4A, B), indicating that OC cells that are highly dependent on SCD also have higher dependency on ATF4/DDIT3. Additionally, expression levels of several important ER stress response genes (*IGFBP1, ERN1, TXNIP, NABP1*) were negatively correlated with SCD (Supplementary Figs. S4C-F), corroborating the association between SCD and ER stress response found through transcriptomic analysis.

### Excess SFAs caused by depletion or pharmacological inhibition of SCD induces ER stress

To further investigate how the imbalance between SFAs and UFAs caused by SCD depletion or inhibition in cancer cells activates the ER stress responses, we next evaluated the two key sensing mechanisms of this pathway: activation of IRE1α and PERK. ER-spanning transmembrane domain of IRE1α and PERK are critical sensors of the levels of lipid saturation in the cell and activate the ER stress response (16). IRE1α causes splicing of the X-box-binding protein 1 (XBP1) mRNA which leads to a spliced form (XBP1s) with potent transcriptional activity (17). PERK autophosphorylates itself upon ER stress and then phosphorylates the eukaryotic translation initiation factor 2α (eIF2α) to halt translation of proteins (18). Translation of ATF4 mRNA, due to its unique 5’ UTR sequence, is upregulated upon phosphorylation of eIF2α (19).

XBP1 splicing was assessed in OVCAR-5 cells stably transduced with shRNA targeting SCD or control shRNA or in cells treated with the SCD inhibitor CAY10566. Increased baseline XBP1s levels were noted in OVCAR-5 and OVCAR-8 cells transduced with shRNA targeting SCD vs control and cultured in low serum medium for 48 hours (Figs. 3A, B & Supplementary Fig. S5A). XBP1 splicing was increased in OVCAR-5 and OVCAR-8 cells transduced with shRNA targeting SCD vs control even under full serum conditions (containing exogenous lipids), but to a lesser degree (Supplementary Figs. S5B, C). Additionally, activation of PERK/CHOP was detected in shSCD vs shCtrl stably transduced OVCAR-8 cells (Supplementary Fig. S5D) under low serum conditions.

**Figure 3.**
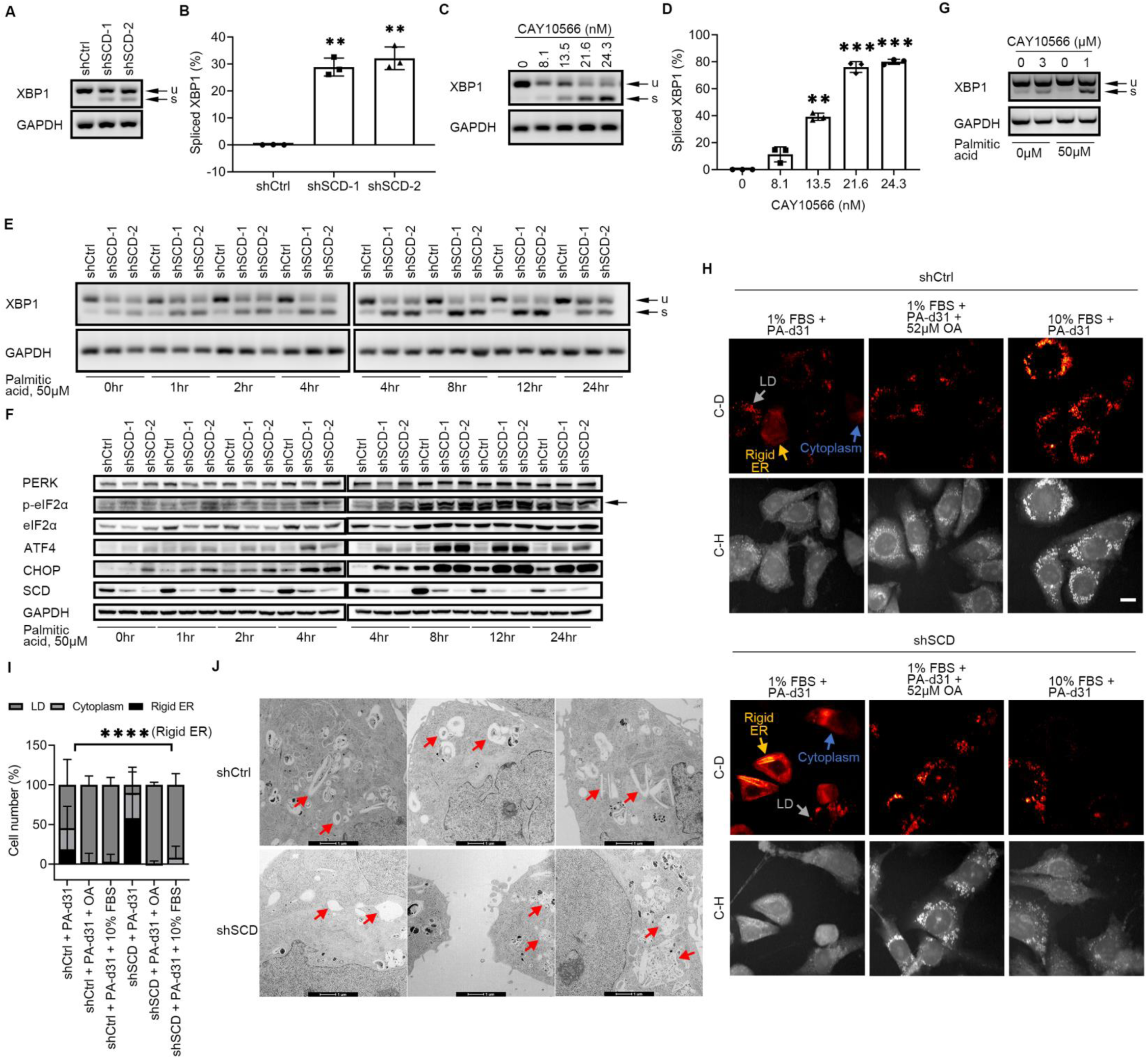
The IRE1α/XBP1 and PERK/eIF2α/ATF4 axes of the ER stress response pathway are activated in ovarian cancer cells under restricted availability of unsaturated fatty acid. (A) XBP1 splicing (u, unspliced transcript; s, spliced transcript) measured by RT-PCR and agarose-gel electrophoresis in OVCAR-5 cells transduced with control shRNA (shCtrl) or shRNAs (1 or 2) targeting SCD (shSCD) and cultured in low serum conditions (1% FBS) for 48 hours. (B) Densitometric analysis of XBP1 splicing products shown in A. Bars represent percent of spliced XBP1 relative to total XBP1 (mean ± SD, n = 3). (C) XBP1 splicing (u, unspliced transcript; s, spliced transcript) in OVCAR-5 cells cultured in low serum conditions and treated with SCD inhibitor CAY10566 for 48 hours. (D) Percent of spliced XBP1 isoform measured by densitometric analysis of PCR products shown in C (mean ± SD, n = 3). (E, F) XBP1 splicing (u, unspliced transcript; s, spliced transcript) (E), and western blot of proteins of the PERK/eIF2α/ATF4 axis (F) in shCtrl and shSCD OVCAR-5 cells cultured under low serum conditions and treated with 50μM palmitic acid for the time periods indicated. (G) XBP1 splicing (u, unspliced transcript; s, spliced transcript) in primary cells from tumors of ovarian cancer patients cultured in low serum conditions and treated with 3μM CAY10566 for 48 hours, or with 1μM CAY10566 and 50μM palmitic acid for 12 hours. (H) Representative SRS images in the C-H and C-D regions of OVCAR-5 shCtrl and shSCD cells cultured in low serum conditions (1% FBS) and treated with 12.5μM palmitic acid-d31 (PA-d31), with or without 52μM oleic acid (OA), or cultured in medium with full serum (10% FBS) and treated with PA-d31 for 24 hr. Yellow arrows indicate rigid ER, gray arrows indicate lipid droplet (LD), and blue arrows indicate cytoplasm. Scale bar: 20 µm. (I) Percentages of shCtrl and shSCD OVCAR-5 cells treated as described in (H) showing C-D SRS signal mainly in rigid ER, lipid droplet (LD), and cytoplasm (n = 139-191). (J) Transmission electron microscopy imaging of smooth ER (red arrows) in OVCAR-5 shCtrl vs shSCD cells cultured in low serum conditions for 48 hours. * p < 0.05, ** p < 0.01, *** p < 0.001, **** p < 0.0001.

Likewise, a dose-dependent increase in XBP1s levels was observed in OVCAR-5 cells treated with CAY10566 (Figs. 3C, D). Having hypothesized that the major trigger of ER stress induced by SCD depletion is the excess of SFAs, XBP1 splicing and activation of PERK/eIF2α/ATF4 were measured in OC cells in which SCD was either knocked down or pharmacologically inhibited in the presence of exogenously added SFA (palmitic acid). We chose a concentration of palmitic acid (50μM) equivalent to that found in media supplemented by 10% FBS(20). Further, as the consequences of ER stress depend on both the length of exposure to cellular stress and the severity of the stress (21), time-dependency was assessed along both the IRE1α/XBP1 and the PERK/eIF2α/ATF4 axes in OC cells supplemented with palmitic acid. The XBP1 splicing signal was increased in OC cells in which SCD was knocked down compared with control cells after supplementation with 50μM of palmitic acid, peaking at approximately 8-12 hours and diminishing at 24 hours (Fig. 3E). Likewise, western blotting analysis of the PERK/eIF2α/ATF4 axis showed increased phosphorylation and corresponding decreased expression levels of eIF2α in OC cells in which SCD was knocked down, starting 2 hours after addition of palmitic acid and peaking at 8-12 hours (Fig. 3F). The downstream transcription factors ATF4 and CHOP were upregulated starting 4 hours after addition of palmitic acid and remaining persistently high up to 24 hours. To confirm these observations in human specimens, we used primary tumor cells dissociated from freshly obtained HGSOC tumors (Supplementary Table S5). Treatment with CAY10566 induced XBP1 splicing in primary HGSOC cells and the XBP1s levels were further augmented by addition of palmitic acid (Fig. 3G & Supplementary Fig. S5E).

Next, to visualize the status of the ER under these conditions, hSRS imaging was performed after treating OVCAR-5 cells stably transduced with control or shRNA targeting SCD with PA-d31 under low serum condition (Fig. 3H). Most of OVCAR-5 shCtrl cells displayed C-D signal derived from PA-d31 in lipid droplets (LD) (depicted as a dots, gray arrow), while a considerable portion of OVCAR-5 shSCD cells displayed the C-D signal on rigid ER (shown as a linear structure, yellow arrow), consistent with an ER stress related morphology (22). Both cell lines contained a small portion of cells harboring detectable C-D signal all over the cytoplasm (Fig. 3H, blue arrow). After PA-d31 treatment alone under low serum conditions, the cell population with rigid ER structures represented the major population of OVCAR-5 shSCD cells, while the cell population with detectable C-D signal in LD was the majority in OVCAR-5 shCtrl cells (Fig. 3H, I). Rescue treatment with UFA (OA or full serum medium) led to the disappearance of rigid ER and increased PA-d31 accumulation in lipid droplets for both shSCD and shCtrl OC cells (Fig. 3H, I), suggesting that supplementation with UFAs can rescue SFA-induced ER stress and facilitate SFA storage in LD. These data support that the balance of intracellular SFA and UFA is essential to prevent ER stress.

Further, examination of OVCAR-5 cells stably transduced with control or shRNA targeting SCD cultured in low serum medium for 48 hours by using transmission electron microscopy (TEM) demonstrated ER membrane disorganization and irregular and compromised ER structure in OVCAR-5 shSCD cells compared to shCtrl cells (Fig. 3J), confirming the biochemical and hSRS imaging results.

### UFAs prevent ER stress in OC cells

To exclude the contribution of potential desaturase-independent functions of SCD to the induction of ER stress in OC cells, we examined whether the effects of SCD depletion or inhibition could be rescued by oleic acid, its main enzymatic product. Supplementation with oleic acid reversed XBP1 splicing in OVCAR-5 cells in which SCD was knocked down (Fig. 4A) as well as in OVCAR-5 cells depleted of SCD and treated with palmitic acid (Fig. 4B). Likewise, supplementation with oleic acid reduced the up-regulation of ATF4 and CHOP along the PERK/eIF2α/ATF4 axis observed in OVCAR-5 cells stably transduced with shRNA targeting SCD and maintained under low serum conditions (Figs. 4C, D). Similar rescue effects of oleic acid on XBP1 splicing were observed in OVCAR-8 and PEO1 cells stably transduced with shRNA targeting SCD (Supplementary Figs. S6A, B).

**Figure 4.**
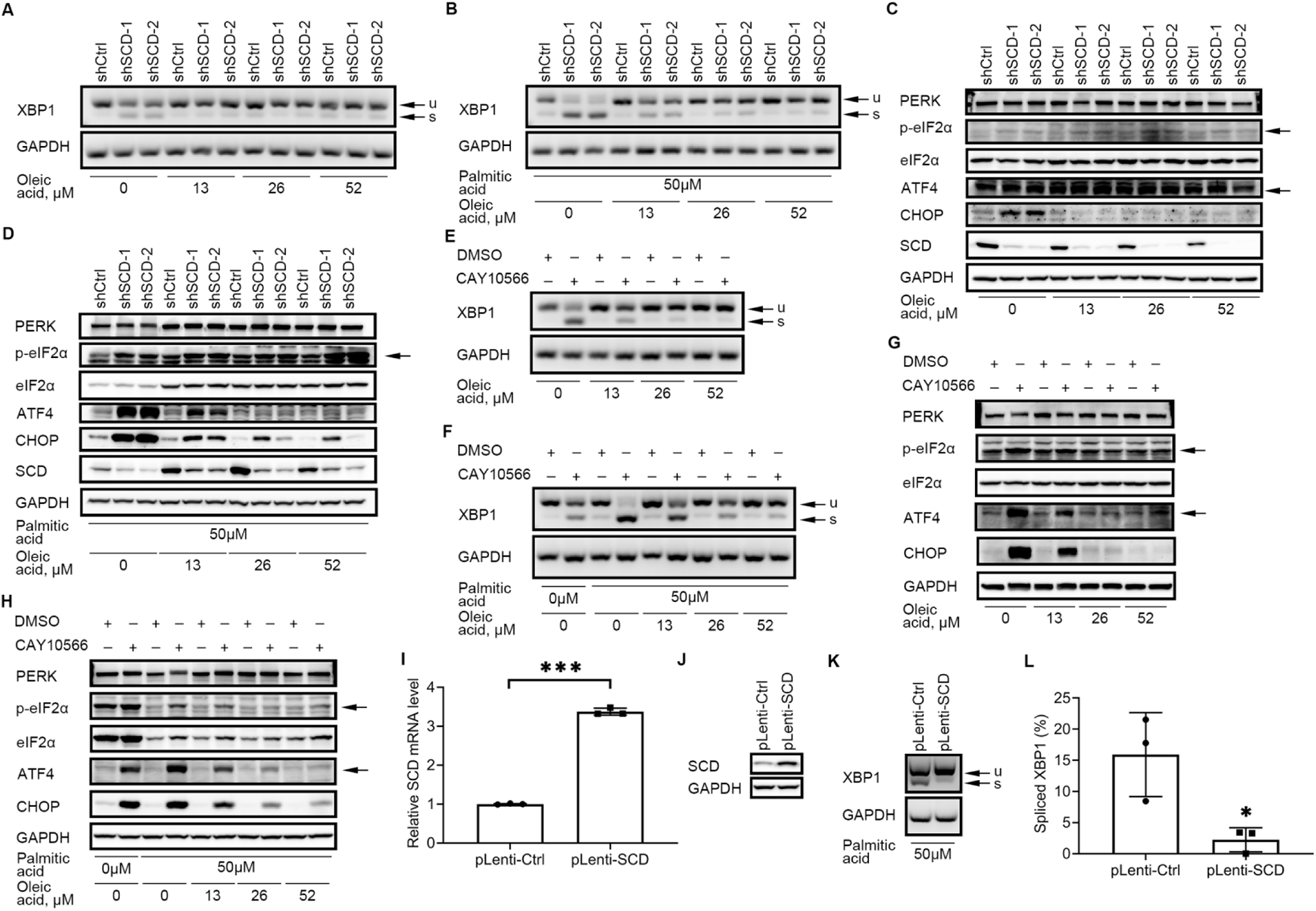
Unsaturated fatty acid-induced ER stress response is reversed by exogenous oleic acid. (A, B) XBP1 splicing (u, unspliced transcript; s, spliced transcript) measured by RT-PCR and agarose gel electrophoresis in OVCAR-5 cells transduced with control shRNA (shCtrl) or shRNAs (1 or 2) targeting SCD (shSCD), cultured in medium containing low serum, and treated with indicated doses of oleic acid for 48 hours (A), or with 50μM palmitic acid and indicated doses of oleic acid for 12 hours (B). (C, D) Western blot of SCD and proteins of the PERK/eIF2α/ATF4 axis in shCtrl and shSCD cells cultured in low serum medium, and treated with different doses of oleic acid for 48 hours (C), or with 50μM palmitic acid and indicated doses of oleic acid for 12 hours (D). (E, F) XBP1 splicing (u, unspliced transcript; s, spliced transcript) in OVCAR-5 cells cultured in medium containing low serum and treated with 21.6nM CAY10566 and indicated doses of oleic acid for 48 hours (E), or with 8.1nM CAY10566, 50μM palmitic acid and indicated doses of oleic acid for 12 hours. (G) Proteins of the PERK/eIF2α/ATF4 pathway measured by western blot in OVCAR-5 cells treated as described in (E). (H) Western blot of proteins of the PERK/eIF2α/ATF4 pathway in OVCAR-5 cells treated as described in (F). Arrows indicate the band of interest. (I, J) Verification of SCD overexpression by qRT-PCR (I) and western blotting (J) in OVCAR-5 cells transduced with a SCD expression vector (pLenti-SCD). Cells transduced with empty vector (pLenti-Ctrl) served as control. Bars represent means ± SD, n = 3. (K) XBP1 splicing (u, unspliced transcript; s, spliced transcript) in pLenti-SCD vs pLenti-Ctrl cells cultured in low serum conditions and treated with 50μM palmitic acid for 12 hours. (M) Percent of spliced isoform (mean ± SD, n = 3) calculated by densitometric analysis of PCR products shown in (L). * p < 0.05, ** p < 0.01, *** p < 0.001, **** p < 0.0001.

Likewise, the rescue effects of oleic acid on ER stress induced by CAY10566 were examined. Supplementation with oleic acid reduced XBP1s levels in OVCAR-5 cells treated with CAY10566 (Fig. 4E), as well as XBP1s levels in cells treated with CAY10566 and palmitic acid (Fig. 4F). Oleic acid also reduced the upregulation of ATF4 and CHOP induced by the SCD inhibitor in OVCAR-5 cells maintained under low serum conditions (Fig. 4G) or in cells additionally treated with 50μM palmitic acid (Fig. 4H).

Lastly, to test the sufficiency of SCD in regulating ER stress triggered by an imbalance of SFA and UFA in OC cells, the enzyme was overexpressed in OVCAR-5 cells. Increased SCD *mRNA* and protein levels were observed in OC cells stably transfected with pLenti-SCD compared to pLenti-Ctrl (Figs. 4I, J). XBP1 splicing induced by addition of palmitic acid in control cells was significantly reduced by SCD overexpression (Fig. 4K, L, p = 0.0277). Collectively, our data support the significance of the balance between SFAs and UFAs regulated by SCD in fine tuning the functions of the ER, likely mediated through direct effects on the ER membrane.

### SCD depletion or inhibition induces apoptosis of OC cells

It is accepted that long-term exposure to mild ER stress or short-term exposure to severe ER stress leads to CHOP-mediated apoptosis (21). RNA-seq analysis of OVCAR-5 SCD KD cells indicated that *DDIT3* (also known as *CHOP, GADD153*) was among the top up-regulated genes in shSCD cells cultured in low serum medium (Fig. 2D, Supplementary Table 4). We therefore hypothesized that OC cells depleted of SCD and cultured under low serum conditions over long period of time would undergo apoptosis. IncuCyte imaging examined Annexin V staining of OVCAR-5 SCD KD cells cultured in low serum medium alone with/without addition of extra palmitic acid in real time. A higher percentage of cells depleted of SCD underwent apoptosis compared with control cells under these conditions. Supplementation with 52μM oleic acid completely rescued the phenotype (Fig. 5A). Further, addition of palmitic acid (50μM) induced early onset of apoptosis in shSCD cells as compared to shCtrl cells, and supplementation with oleic acid reversed the phenotype (Fig. 5B). Likewise, CAY10566 induced apoptosis in OVCAR-5 cells grown under low serum conditions, starting at 48 hours; while oleic acid supplementation blocked this effect (Fig. 5C). Addition of palmitic acid to the CAY10566 treatment caused earlier onset of apoptosis (less than 24 hours) and augmented the phenotype (>20% apoptotic cells; Fig. 5D). Similarly, oleic acid supplementation rescued the phenotype (Figure 5D). On the other hand, stable SCD knock down did not affect proliferation of OVCAR-5 cells compared to controls when cells were maintained under full serum conditions. However, SCD KD decreased proliferation of cells cultured under low serum conditions (Supplementary Figs. S6C, D).

**Figure 5.**
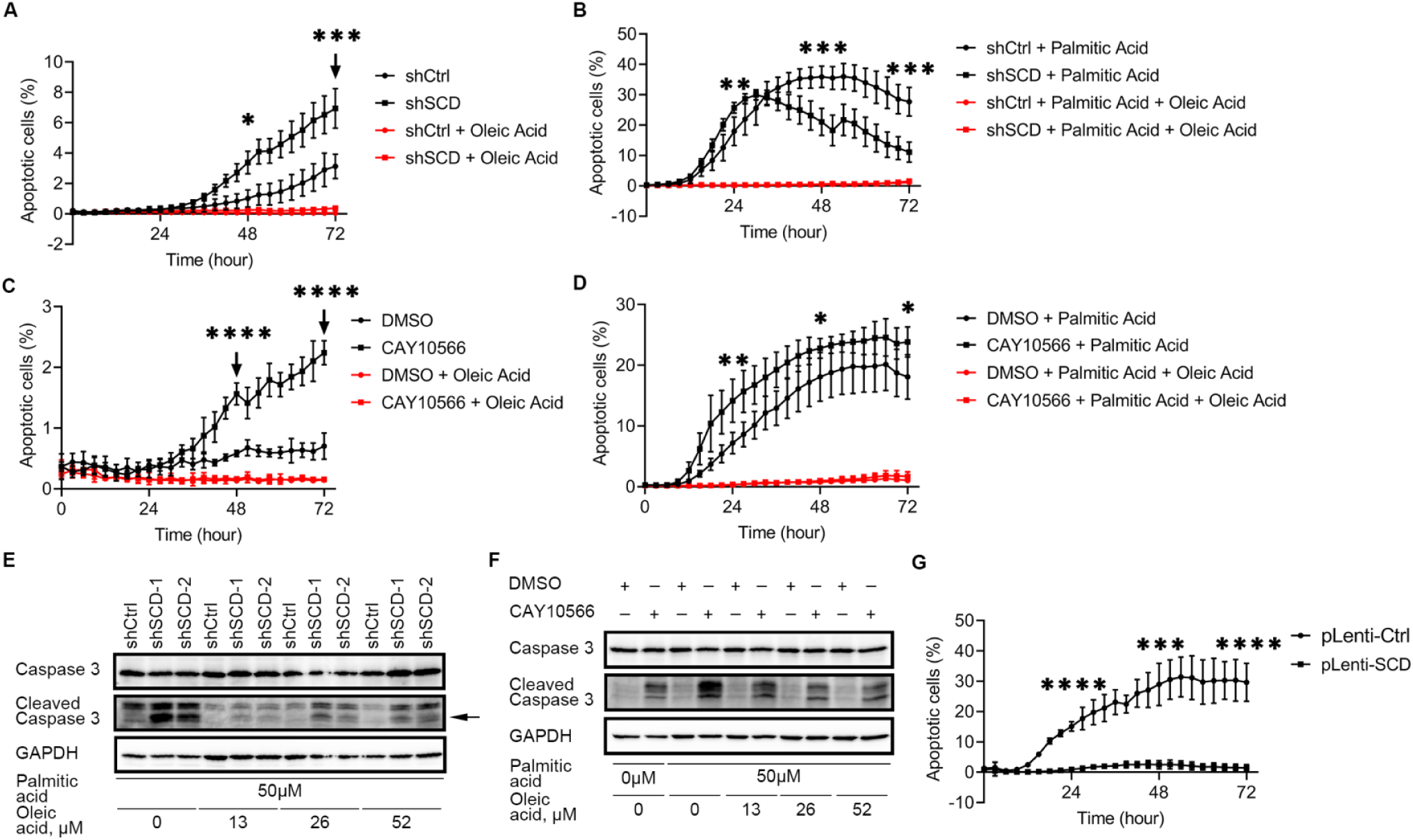
Apoptosis induced by SCD inhibition, or treatment with palmitic acid, is attenuated by exogenous oleic acid or SCD overexpression. (A-B) Time-lapse imaging of Annexin V staining to measure apoptosis in OVCAR-5 cells transduced with control shRNA (shCtrl) or shRNA targeting SCD (shSCD), cultured in low serum conditions, and treated with 52μM oleic acid (A), or with 50μM palmitic acid alone or in combination with 52μM oleic acid (B). (C-D) Time-lapse of Annexin V imaging to measure apoptosis in OVCAR-5 cells cultured in low serum conditions and treated with 21.6nM CAY10566, 52μM oleic acid or combination (C), or with 8.1nM CAY10566, 50μM palmitic acid, 52μM oleic acid, or combinations (D). (E) Western blot of full-length and cleaved caspase-3 in shSCD vs shCtrl cells cultured under low serum medium and treated with 50μM palmitic acid and indicated doses of oleic acid for 12 hours. (F) Western blot of full-length and cleaved caspase-3 in OVCAR-5 cells cultured in low serum medium and treated with 8.1nM CAY10566, 50μM palmitic acid and indicated doses of oleic acid for 12 hours. (G) Time-lapse of Annexin V imaging to determine apoptosis in OVCAR-5 cells overexpressing SCD (pLenti-SCD) and control cells (pLenti-Ctrl), cultured in medium containing low serum and treated with 50μM palmitic acid. Values in panels A-D & G are means ± SD, n = 6. * p < 0.05, ** p < 0.01, *** p < 0.001, **** p < 0.0001.

To confirm the apoptosis phenotype, we also examined the cleavage of caspase-3 in OVCAR-5 transduced with shRNA targeting SCD vs. control shRNA cultured under low serum conditions and treated with extra palmitic acid. Increased caspase-3 cleavage was observed in OVCAR-5 shSCD compared to shCtrl cells (Fig. 5E). Importantly, cleavage of caspase-3 in cells depleted of SCD and treated with palmitic acid could be reversed by supplementation with oleic acid (Fig. 5E). Caspase-3 cleavage was also induced by CAY10566, augmented by addition of extra palmitic acid and rescued by restoration of the SFA/UFA balance, through repletion of oleic acid (Fig. 5F).

Lastly, to test the sufficiency of SCD to this phenotype, OVCAR-5 cells in which SCD was overexpressed (pLenti-SCD) vs control cells (pLenti-Ctrl) were maintained under low serum conditions and were treated with 50μM palmitic acid for 72 hours. The percentage of apoptotic cells induced by exogenous palmitic acid was significantly reduced in cells in which SCD was overexpressed as compared to control cells (Fig. 5G), supporting that OC cells with higher SCD expression could survive the stress induced by SFA. Together, the data demonstrate that the balance between intracellular UFA and SFA regulated by the desaturase SCD is critical to determining the survival of OC cells.

### SCD depletion or inhibition suppresses tumor growth *in vivo*

Whether the observed effects of SCD on cancer cell survival impact tumor progression *in vivo* remains unknown. Tumorigenicity of OC cells depleted of SCD was tested by using an intraperitoneal (ip) xenograft model in athymic nude mice. SCD knockdown compared to control OVCAR-5 cells led to significantly reduced tumor weight (234.3 mg vs. 360.1 mg, p = 0.0343) and tumor volume (209.2 mm^3^ vs. 404.8 mm^3^, p = 0.0212; Figs. 6A, B), although the numbers of peritoneal metastases (Fig 6C, 208.8 vs. 217.3, p = 0.7602) and ascites volume (Supplementary Fig. S7A) were similar between shSCD and control groups. SCD knock down was maintained *in vivo*, as demonstrated by qRT-PCR measurement of *SCD* in RNA extracted from harvested tumors (Fig. 6D, p value = 0.0015). Furthermore, increased XBP1s levels were observed in xenografts derived from OVCAR-5 cells stably transduced with shRNA targeting SCD compared to controls (Figs. 6E, F, p = 0.0018), supporting that an increased susceptibility to ER stress is maintained *in vivo* in tumors depleted of the enzyme. Similar inhibitory effects on tumor growth were observed in a subcutaneous mouse xenograft model derived from OVCAR-3 cells stably transduced with shRNA targeting SCD vs. shRNA control (Supplementary Figs. S7B, C).

**Figure 6.**
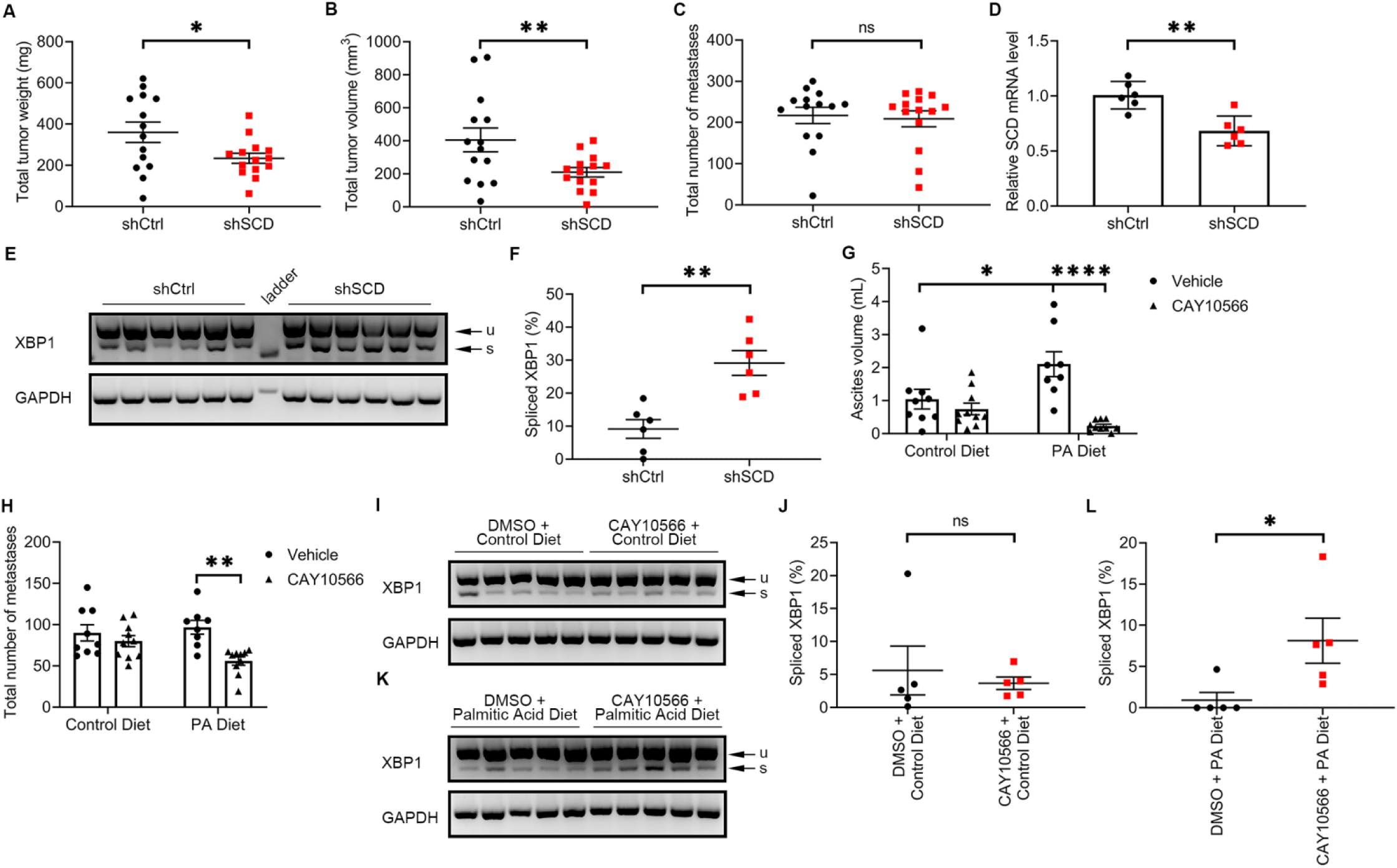
SCD knockdown inhibited growth of ovarian cancer xenografts in mice. (A-C) Total tumor weight (A), total tumor volume (B) and total number of metastases (C) in athymic nude mice intraperitoneally injected with OVCAR-5 cells transduced with control shRNA (shCtrl) or shRNA targeting SCD (shSCD), and evaluated after 28 days (values are means ± SE, n = 14 per group). (D) qRT-PCR measurements of SCD expression (mean ± SD, n = 6) in a random sample of tumor xenografts described in (A). (E, F) Agarose gel electrophoresis of XBP1 splicing products (u, unspliced transcript; s, spliced transcript) (E), and percent spliced XBP1 isoform estimated by image analysis of transcript bands (mean ± SD, n = 6) (F) in a random sample of tumor xenografts described in (A). (G, H) Ascites volume (G) and total number of metastases (H) in athymic nude mice intraperitoneally injected with OVCAR-5 cells, fed with a palmitic acid-rich diet or control diet, and treated with SCD inhibitor CAY10566 or vehicle for 28 days. Values are means ± SE, n = 10. (I-L) XBP1 splicing products (u, unspliced transcript; s, spliced transcript) (I, K), and percent intensity of the spliced transcript estimated by image analysis (J, L) in a random sample (n = 5) of tumor xenografts described in (F). Values are means ± SD. * p < 0.05, ** p < 0.01, *** p < 0.001, **** p < 0.0001.

Our observations that ER stress and apoptosis are augmented *in vitro* by inhibition of the desaturase coupled with an excess of SFAs led us to hypothesize that the effects of a pharmacological inhibitor targeting this pathway would be enhanced through dietary modifications tilting the FA balance towards increased saturation levels. To test this hypothesis, we used an OC ip model fed a palmitic acid rich or control diet, and treatment with CAY10566 or diluent. While CAY10566 did not significantly alter the numbers of peritoneal metastases and the ascites volume in mice fed a control diet (Figs. 6G, H), the inhibitor significantly decreased both the ascites volume (p < 0.0001) as well as the numbers of metastases (p = 0.0003) in mice fed palmitic acid rich diet (Figs. 6G, H). Interestingly, an unanticipated, but significant increase in ascites volume was observed in the group receiving the SFA rich diet compared with the control diet (Fig. 6G, p = 0.0252). No significant differences were observed in total tumor weights and total tumor volumes (Supplementary Figs. S8D, E). XBP1s levels were not significantly different in tumors derived from mice fed the control diet and treated with vehicle vs CAY10566 (Figs. 6I, J, p = 0.6370), but were significantly increased in tumors derived from mice fed palmitic acid rich diet and treated with CAY10566 vs. control (Figs. 6K, L, p= 0.0369). The data support that inhibition or depletion of SCD exerts anti-tumorigenic effects through a mechanism dependent on ER stress induction likely caused by decreased levels of lipid desaturation.

## Discussion

Our data based on transcriptomic, lipidomic analyses, SRS and EM imaging demonstrate that the imbalance between saturated and unsaturated fatty acids caused by depletion or inhibition of the enzyme SCD has a key function in determining cancer cell survival or death and impacts tumor progression *in vivo*. These findings hone on a largely understudied metabolic process in cancer cells and suggest the possibility of using an intervention combining an SCD enzymatic inhibitor together with a dietary intervention to increase levels of lipid saturation in cancer cells and trigger a potent anti-tumor effect. Our findings have several important consequences.

First, our results point to the significance of the process of lipid unsaturation in cancer. Lipid synthesis includes the synthesis of FA from acetyl-coA mediated by fatty acid synthase (FASN) and acyl-CoA synthase (ACSL), followed by addition of double carbon bonds catalyzed by FA desaturases. Among the three desaturases, Δ9 (stearoyl-CoA desaturase-1, SCD) catalyzes the synthesis of monounsaturated fatty acids by adding one double bond to saturated fatty acids (mostly stearic acid), while the Δ6 and Δ5 desaturases are involved predominantly in the synthesis of polyunsaturated fatty acids (23). The current study builds on our previous observations that highly tumorigenic ovarian CSCs are enriched in UFAs, caused by SCD upregulation, which in turn, induce pro-survival NF-κB signaling allowing CSCs to proliferate as spheres and effectively initiate tumors in immunodeficient mice (11). Here we show that beyond the previously observed effects on CSCs, SCD knockdown, and to a lesser extent its inhibition, induces anti-tumor effects in intraperitoneal xenografts which closely resemble the pattern of dissemination of the human disease. Overexpression of ACSL and SCD leading to increased UFA levels was described in association with epithelial to mesenchymal transition and aggressive clinical prognosis in colon cancer, and SCD upregulation was observed in basal type breast cancer and aggressive prostate cancer (24, 25). Depletion of oleic acid (but not of palmitate) inhibited the proliferation of acute meylogenous leukemia and lymphoma cells (26). Unsaturated lipids also play a role in maintaining the membrane fluidity and support signaling through membrane receptor tyrosine kinases (27). Lipid saturation detectable in plasma or in tumor tissue was associated with cancer risk in several epidemiological studies (28, 29). For example, decreased risk of breast cancer was noted in women with high plasma stearic acid levels and low SCD activity, supporting that high lipid saturation is indirectly correlated with cancer risk (29). However, it remains not clear how the balance between UFAs and SFAs alter cancer predisposition or cancer progression, a concept we addressed here.

A powerful tool that allowed us to quantify UFAs and SFAs at the single cell level in this study was SRS microscopy, an innovative label-free chemical imaging technique which we have previously employed to characterize metabolic reprograming from glycolysis to fatty acid uptake and β-oxidation in platinum-resistant cancer cells (30), increased lipid desaturation in ovarian CSCs (11), and cholesteryl ester accumulation as a metabolic marker for multiple aggressive cancers (31, 32). Combined with post-processing methods such as least-square fitting, multivariate curve resolution and phasor segmentation, hSRS can distinguish different intracellular biomolecules such as protein and fatty acid (10, 33). Here, SRS imaging visualized SFA-induced ER stress in OC cells depleted of SCD and the rescue observed by manipulating the ratio between SFAs and UFAs. We recognize that highly spatially overlapping biomolecules with similar spectrum profile such as SFAs and UFAs are difficult to decompose by using conventional analysis methods. We propose that pixel-wise LASSO unmixing can resolve such shortcomings, enabling the simultaneous evaluation of multiple metabolites, facilitating better assessment of cancer cell metabolism and metabolic changes in response to gene modification and drug treatment. The innovative hSRS-LASSO imaging method utilized here provides a new way to explore cancer lipid metabolism in deeper detail.

Our findings demonstrate that the imbalance between SFAs and UFAs impedes cancer cell survival and *in vivo* tumor growth through induction of ER stress. We observed that two branches of ER stress response, regulated by the PERK and IRE1α were activated in response to SCD depletion or inhibition, that this activation was augmented in the presence of SFA supplementation and could be rescued by addition of UFAs. Interestingly, non-cancer cells, such as neural stem/progenitor cells were found to be susceptible to lipid accumulation and consequent ER stress (34) and palmitic acid was shown to induce ER stress and apoptosis in human bone marrow-derived mesenchymal stem cells(35). Other recent studies found that N-Myc driven hepatocellular cancer cell or glioma proliferation was dependent on lipid unsaturation through modulation of ER stress responses (36). While a definitive mechanistic explanation is still lacking, our hSRS and TEM imaging suggest that increased levels of lipid unsaturation in cancer cells alter the morphology of the ER, causing increased membrane rigidity and potentially direct activation of the sensor proteins.

The link between ER stress and tumor progression is emerging. In orthotopic cervical cancer xenografts, hypoxia-induced metastasis to lymph nodes was blocked by PERK inhibition (37). The PERK/eIF2α/ATF4 axis was also shown to be important for inducing epithelial-to-mesenchymal transition of breast cancer cells (38), while the IRE1α/XBP1 pathway was involved in tumor initiation and breast cancer progression by activating the transcription factor HIF1α (39). Our study, together with previous reports, reveals the biphasic role of PERK/eIF2α/ATF4 axis of the ER stress response; with the initial response being implicated in salvaging the stressed cells, whereas prolonged activation of the pathway leads to apoptosis.

Future investigation of how lipid unsaturation triggers ER stress and modulates the ER stress response could provide new therapeutic opportunities to target cancer progression.

A starting point towards this goal is our attempt to combine SCD inhibition with a dietary intervention. We demonstrate that this intervention resulting in accumulation of toxic SFAs has potent anti-tumor effects and induces ER stress responses in tumors. The putative role of diet in cancer treatment has remained a “holy grail” and is based on the concept that altered metabolite levels in the tumor microenvironment could create metabolic challenges impeding tumor cells’ survival. A recent study found that severe caloric restriction led to inhibited tumor growth (40) and a proposed mechanism, was the downregulation of SCD, resulting in an imbalance between SFAs and UFAs, similar to what we achieved in our study through SCD pharmacological inhibition and supplementation with palmitic acid. The role of saturated lipids in cancer remains controversial, with the overwhelming belief that saturated fat is pro-tumorigenic. Indeed, a recent study showed that dietary palmitic acid promoted metastasis in oral carcinoma and melanoma models (41). Specifically, *in vitro* palmitic acid treatment primed tumor cells for metastasis after implantation *in vivo*, while a palmitic acid-rich diet promoted increased metastatic tumor burden. Consistent with these observations, we found that the palmitic acid-rich diet induced increased accumulation of malignant ascites, the main vehicle of OC intraperitoneal metastasis. However, combined palmitic acid-rich diet and the SCD inhibitor CAY10566, significantly reduced the volume of ascites as well as the numbers of metastases, with signals of ER stress response being detectable directly in tumor tissue.

Given the plasticity and redundancy of metabolic pathways, which have by and large thwarted prior attempts to use metabolic interventions to target cancer, our results highlight the importance of combination strategies. Taken together our findings support SCD’s key role in regulating cancer cell fate and tumorigenicity and point to new modalities to block OC progression.

## Materials and Methods

### Cell culture

OVCAR-5 cells and OVCAR-8 cells were provided by Dr. Marcus Peter, Northwestern University, and Dr. Kenneth Nephew, Indiana University, respectively. PEO1 cells were purchased from MilliporeSigma (cat#: 10032308). OVCAR-3 cells were purchased from the American Type Culture Collection (ATCC) (cat#: HTB-161). FT-190 cells (immortalized human fallopian tube luminal epithelial cells) were provided by Dr. Ronny Drapkin, University of Pennsylvania. OVCAR-5, PEO1 and OVCAR-3 cells were cultured in RPMI-1640 with L-glutamine (Corning cat#: 10-040-CV), OVCAR-8 cells were cultured in DMEM (Corning cat#: 10-017-CV). Media was supplemented with 10% FBS (Corning cat#: 35011CV), 1% GlutaMAX (Gibco cat#: 35050-061), and 100μg/mL penicillin/streptomycin (Cytiva cat#: SV30010) and all cells were maintained at 37°C in an incubator with 5% CO_2_ and 100% humidity. For *in vitro* experiments, cells were cultured in low serum medium (1% FBS) unless otherwise stated. Treatment of 50μM palmitic acid (MilliporeSigma cat#: P0500) recapitulated the concentration of palmitic acid in 10% FBS media (20). Oleic acid (MilliporeSigma cat#: O3008) was added at different doses with the lowest dose of 13μM equivalent to that in media containing 10% FBS (20). All cell lines were authenticated by IDEXX BioAnalytics and were determined to be free of mycoplasma contamination by IDEXX BioAnalytics or Charles River Laboratories. In addition, cells were regularly tested for mycoplasma in our laboratory using a Universal Mycoplasma Detection Kit (ATCC cat#: 30-1012K).

### Reagents

Palmitic acid (cat#: P0500), oleic acid (cat#: O3008), dimethyl sulphoxide (DMSO) (cat#: D2650) and Tween® 80 (cat#: P4780) were purchased from MilliporeSigma. CAY10566 (cat#: HY-15823) and PEG300 (cat#: HY-Y0873) were purchased from MedChemExpress. Saline (cat#: Z1376) was purchased from Intermountain Life Sciences.

### Human specimens

An ovarian cancer tissue microarray (TMA) was obtained from the Cooperative Human Tissue Network (CHTN; OvCa2), which contains different histological subtypes of ovarian cancer, benign ovarian tumors, and normal fallopian tube (see SM). Primary tumors or ascites from high grade serous ovarian cancer patients were collected at Northwestern Memorial Hospital from consenting donors. Tumor tissues were minced into small pieces and digested with in DMEM/F-12 medium (Gibco cat#: 11320-033) supplemented with 300U/mL collagenase (MilliporeSigma cat#: C7657) and 300U/mL hyaluronidase (MilliporeSigma cat#: H3506) at room temperature overnight. The next day, tissues were digested with trypsin (Corning cat#: 25054CI) at 37°C for 10min, followed by treatment with 1X red blood cell lysis buffer (Biolegend cat#: 420301) on ice for 10min, and then with DNase I at 37°C for 10min to produce a single-cell suspension. Cells were resuspended in RPMI-1640 with L-glutamine supplemented with 10% FBS, 1% GlutaMAX, and 1% Pen/Strep. For ascites, cells were spun down and resuspended in RPMI-1640 as described above.

### Cell transfection; construction of SCD expression vector, RNA extraction and real-time qPCR, western blot and immunohistochemistry (IHC)

were performed using protocols described previously (42-45) and detailed in the Supplementary Material (SM). Primers are included in Supplementary Table S1.

### XBP1 splicing assay

Synthesized cDNA as described above was used to measure levels of unspliced and spliced XBP1mRNA were measured by regular PCR performed with GoTaq Green Master Mix (Promega cat#: M7123) on a T100 Thermo Cycler (Bio-Rad cat#: 186-1096). The primers targeting the spliced XBP1 region (116bp) (Supplementary Table S1) were designed according to a previous study (46). PCR products were resolved by agarose gel electrophoresis and visualized using GelGreen stain (Biotium cat#: 41005) on an ImageQuant LAS 4000 machine (GE Healthcare). Densitometric analysis of product bands was performed using the Gel Analyzer function in Fiji from NIH (47).

### Lipidomics

Cell pellets were obtained from OVCAR-5 cells transduced with shRNA targeting SCD vs. control shRNA cells used for analysis at the Bindley Bioscience Center, Purdue University. Lipidomic analysis was performed using multiple reaction monitoring (MRM) profiling, as detailed in SM. For reporting of the relative amounts using normalization by the internal standards, the amount of each fatty acid was expressed as pg/1000 cells. For lipidomics profiling using the relative amounts, cells were lysed in ultrapure water for lipid extraction. Statistical analysis was performed utilizing MetaboAnalystR 3.0 (48). Data on the relative amounts from different lipid classes were scaled to obtain a normal distribution, and evaluated by univariate analysis, principal component analysis (PCA), and cluster analysis/heatmap. Informative lipids were analyzed according to class, fatty acyl residue chain unsaturation level.

### SRS imaging

Stimulated Raman scattering (SRS) imaging was performed to measure isotope labelled cellular saturated/unsaturated fatty acids on a previously described lab-built system with a femtosecond laser source operating at 80MHz (InSight DeepSee, Spectra-Physics, Santa Clara, CA, USA) (11, 30). Briefly, the laser source provides two synchronized output beams, a tunable pump beam ranging from 680 nm to 1300 nm and a fixed 1040 nm Stokes beam, modulated at 2.3MHz by an acousto-optic modulator (1205-C, Isomet). SRS spectrum is obtained by controlling the temporal delay of two chirped femtosecond pulse. A 12.7cm long SF57 glass rod was used to chirped Stoke path to compensate for its longer wavelength. After combination, the path of both beams was further chirped by five 12.7cm long SF57 glass rods before sent to a laser-scanning microscope. A 60x water immersion objective (NA = 1.2, UPlanApo/IR, Olympus) was used to focus the light on the sample, followed by signal collection via an oil condenser (NA = 1.4, U-AAC, Olympus). For hyperspectral SRS (hSRS) imaging, a stack of 120 images was recorded at various pump-Stokes temporal delay, implemented by tuning the optical path difference between pump and Stokes beam through a translation delay stage. Pump beam was tuned to 798 nm for imaging at the C-H vibration region (2800 ∼ 3050 cm^-1^), and to 850 nm for imaging at C-D vibration region (2100 ∼ 2300 cm^-1^). The power of pump and Stokes beam before microscope was 30mW and 200mW respectively. Raman shift was calibrated by standard samples, including DMSO, DMSO-d6, palmitic acid-d31 (PA-d31) and oleic acid-d34 (OA-d34). Hyperspectral SRS images were analyzed using ImageJ and previously described pixel-wise least absolute shrinkage and selection operator (LASSO) regression algorithm (49). In brief, LASSO can effectively decompose hSRS imaging into maps of different biomolecules by introducing a sparsity constraint to suppress the crosstalk between different chemical maps. PA-d31 and OA-d34 was used to provide reference spectrum of SFA and USFA respectively for LASSO unmixing analysis. To study cellular uptake of fatty acids by SRS microscopy, cells were seeded on 35 mm glass-bottom dishes (Cellvis, D35-20-1.5-N) overnight, and then cultured in low serum medium for 24 hours, followed by treatment with 12.5μM PA-d31 (Cambridge Isotope Lab) for 24 hours. Rescue experiments were conducted by adding 52μM oleic acid to the low serum medium containing 12.5μM PA-d31 or changing to full serum medium with 12.5μM PA-d31. For quantitative SRS imaging, cells were fixed with 10% neutral buffered formalin for 30 min followed by 3 times of PBS wash.

### Xenograft experiments

Athymic nude mice (strain: *Foxn1*^*nu*^), 6-8 weeks old, were obtained from Envigo (Indianapolis, IN, USA). For a subcutaneous model, mice were injected with 2×10^6^ OVCAR-3 cells stably transduced with shRNAs targeting SCD or control shRNA resuspended in 1/1 mix of RPMI 1640 basal medium and Matrigel (Corning cat#: 356234). Tumor length (L), width (W) and height (H) were measured every three days with digital calipers. Tumor volume was calculated using the formula V = L × H × W/2. For an intraperitoneal model, 5 million cells OVCAR-5 cells, or cells stably transduced with shRNAs targeting SCD or control shRNA were injected intraperitoneally (ip). Mice were euthanized 28 days after injection of cells to evaluate tumor growth and abdominal ascites. To determine the effects of a palmitic acid-enriched diet and SCD inhibitor combination, athymic nude mice were randomized to be fed with a palmitic acid-enriched or a control diet beginning one week before ip injection of OVCAR-5 cells, and continued for the duration of the experiment (28 days). Mice were treated ip with the SCD inhibitor CAY10566 (8 mg/kg body weight) or diluent (10% DMSO, 40% PEG300, 5% Tween-80, 45% saline), every other day during weekdays. The contents of palmitic acid in the palmitic acid-enriched diet and control were 30% and 11.7% of the total fatty acids, respectively. The levels of total fat (58g/kg) were the same, and levels of saturated fat, monounsaturated fat, polyunsaturated fat, stearic acid, and oleic acid were similar (< 2% difference) between the palmitic acid-enriched and control diets. The diets were custom-made by Teklad (Envigo).

### RNA sequencing (RNA-seq)

was performed as previously described (42, 43) and detailed in SM. Differentially expressed (DE) genes between experimental groups were determined and FDR corrected for multiple hypothesis testing with the edgeR package (50) in R. Pathway analysis based on the differentially expressed genes was performed using Ingenuity Pathway Analysis (IPA) software (QIAGEN).

### The Genotype-Tissue Expression (GTEx) and The Cancer Genome Atlas (TCGA) ovarian cancer RNA-seq data analysis

Recomputed RSEM expected counts of fallopian tube samples (GTEx) and primary tumor samples from TCGA ovarian cancer patients were downloaded from UCSC Xena Browser. Expression of SCD (Ensembl ID: ENSG00000099194) was extracted from matrix of all genes and Student’s t test was performed on log2 transformed RSEM expected counts between fallopian tube samples and primary tumor samples. Data are deposited in GEO (GSE192442).

### Cancer Dependency Map data analysis

Dependency score data from CRISPR screening in cancer cell lines was downloaded from Depmap Portal (14). Ovarian cancer cell lines were selected and Pearson correlation score was calculated between SCD and transcriptional regulators identified from IPA.

### Apoptosis assay by IncuCyte imaging

Cells were seeded on 96-well plates at 500 per well and cultured in an IncuCyte S3 live-cell analysis system. Serum was reduced in the medium to 1% FBS, and Annexin V Green Dye (Sartorius cat#: 4642) was added for detection of apoptotic cells under different experimental conditions. Four images of bright field and green fluorescence were captured at three-hour intervals during a 72hr evaluation period. Cells were counted on the images and expressed as percentage of apoptotic cells (green cells/total cells under bright field).

### Transmission Electron Microscopy

Cells were seeded at a density of 32,000 on glass bottom dishes (Cellvis cat#: D35-14-1.5-N) and cultured in regular medium for 48 hours. Culture conditions were changed to low serum medium (1% FBS) for an additional 48 hours. Cells were rinsed twice with PBS and fixed with 0.1M sodium cacodylate buffer, pH 7.3, containing 2% paraformaldehyde and 2.5% glutaraldehyde. Cells were post-fixed with 2% osmium tetroxide in unbuffered aqueous solution followed by rinsing with distilled water. Subsequently, cells were *en-bloc* stained with 3% uranyl acetate and rinsed with distilled water. Finally, cells were dehydrated in ascending grades of ethanol, transitioned with 1:1 mixture of ethanol and resin, and embedded in resin mixture of EMbed 812 Kit (Electron Microscopy Sciences cat#: 14120), cured in a 60°C oven. Samples were sectioned on a Leica Ultracut UC6 ultramicrotome. Sections (70 nm) were collected on 200mesh copper grids and post stained with 3% uranyl acetate and Reynolds lead citrate. Electron microscopic images were captured on a FEI Tecnai Spirit G2 transmission electron microscope.

## Statistical Analyses

Data were analyzed by two-tailed Student t test. Two-way ANOVA was used for the analyses of the *in vivo* experiment with CAY10566 and palmitic acid-enriched diet. IHC staining analysis employed the Fisher exact test to compare the groups. Tumor weights and volumes from the subcutaneous mouse xenograft experiment were log transformed as paired samples before statistical analysis. P < 0.05 was considered statistically significant. All statistical analyses except for RNA-seq and lipidomics were performed using GraphPad Prism v8.0.

## Data Availability

The RNA-sequencing data generated during the current study are available in the Gene Expression Omnibus repository with accession ID: GSE192442. OVCAR-5 SCD KD lipidomics profiling significant lipids species list and OVCAR-5 SCD KD RNA-seq significant genes list were included in the Supplementary Information Appendix. The datasets of all lipid species from untargeted lipidomics experiment during the current study are available from the corresponding author on reasonable request.

## Acknowledgements

This work was supported by R01 CA224275 to DM and JXC, R33 CA223581 and NSF CHE1807106 grants to JXC. We thank Drs. Debabrata Chakravarti and Navdeep Chandel for valuable comments. Tumor specimens were procured through the Pathology Core and sequencing was performed in the NUSeq Core supported by NCI CCSG P30 CA060553 awarded to the Robert H Lurie Comprehensive Cancer Center. Lipidomics analysis was performed in Metabolite Profiling Facility at Bindley Bioscience Center, Purdue University. Transmission electron microscopy was performed in the Center for Advanced Microscopy/Nikon Imaging Center (CAM) at Northwestern University supported by NCI CA060553. This research was supported in part through the computational resources and staff contributions provided for the Quest high performance computing facility at Northwestern University which is jointly supported by the Office of the Provost, the Office for Research, and Northwestern University Information Technology.

## Author Contributions

JXC and DM designed this study. GZ, YT and DM drafted the manuscript. GZ, YT, HC, DV and JJW performed the experiments and collected the data. GZ, YT, HC, HH and YW conducted the statistical analysis. CRF performed the lipidomics experiment and initial analysis. RK contributed to TCGA and GTEx data analysis. HC, JXC and DM revised this manuscript. All authors read and approved the final manuscript.

### Ethics declarations

Human subject studies were conducted in accordance with the Declaration of Helsinki and approved by the IRB (Northwestern University IRB#: STU00202468). Animal studies were conducted according to a protocol approved by the Institutional Animal Care and Use Committee of Northwestern University (# IS00008973).

