## Supplementary Material for "The Balance between Saturated and Unsaturated Fatty Acids Regulates Ovarian Cancer Cell Fate"

**Cell Transduction.** Gene knock down was performed by plating cells at a density of 40,000 per well in a 24-well plate (day 0). On day 1, cells were transduced with lentiviral particles containing shRNAs targeting human SCD (MilliporeSigma cat#: SHCLNV) or with scrambled (control) shRNAs (MilliporeSigma cat#: SHC001V). Lentiviral particles were diluted in complete cell culture medium containing 8μg/mL polybrene to obtain a multiplicity of infection (MOI) of 5. Cells were incubated with lentiviruses for 24 hours, medium containing lentiviruses was removed, and then cells were cultured in complete medium for another 24 hours. On day 3, complete medium containing 2.0μg/mL puromycin (Gibco cat#: A1113803) was added to cell cultures to start selection of transfected cells. After selection (1-2 weeks), cells were passaged 3-4 times before gene knock down was verified in the stable cells by real-time RT-PCR.

**Construction of SCD expression vector.** The *SCD* coding sequence was amplified by PCR from a SCD Human Tagged ORF Clone plasmid (OriGene cat#: RC209148) using the following primers 5'-GCTCTAGA GGATCCACCGGTCGCCACCATGCCGGCCCACTTG-3' (forward) and 5'- GACGTCGACGCGGCCGCTTCAGCCACTCTTGTAGTTTCCATC-3' (reverse), and then inserted into the pLenti-CMV-Neo vector (Addgene plasmid#: 17447). Lentiviral particles were packaged in HEK-293T cells via co-transfection with pMD2.G (Addgene plasmid#: 12259) and psPAX2 (Addgene plasmid#: 12260). Viral particles were harvested 48hrs and 72hrs post transfection, pooled, and concentrated using Lenti-X Concentrator (TaKaRa cat#: 631231). Pellets of lentiviral particles were resuspended in PBS (Cytiva cat#: SH30256.LS), aliquoted, and stored at -80℃.

**RNA extraction and real-time qPCR.** Total cellular RNA was extracted using TRI reagent according to the manufacturer's instruction (MilliporeSigma cat#:T9424). cDNA was synthesized from 1µg of total RNA with a High Capacity cDNA Reverse Transcription Kit (Applied Biosystems cat#: 4368814). Genes of interest were amplified by PCR using PowerUp SYBR Green Master Mix (Applied Biosystems cat#: A25742) on a QuantStudio 5 Real-Time PCR System (Thermo Fisher Scientific cat#: A28140) and quantified applying the $2^{-\Delta\Delta CT}$ method (2) using GAPDH for normalization. Primers used for real-time qPCR are described in Supplementary Table S1.

**Western blot.** Cell lysates were prepared using Radio-immunoprecipitation assay (RIPA) buffer. Protein concentrations were measured with the Bradford reagent (Bio-Rad Protein Assay, cat#: 5000006) using BSA as standard. Protein lysates (20µg per sample) were denatured at 100°C, resolved by polyacrylamide-gel electrophoresis, and transferred onto a PVDF membrane. The membrane was incubated with 5% milk (blocking), and then with primary antibody overnight at 4℃, followed by incubation with HRP-conjugated secondary antibody for 1hr at room temperature. Signal was developed using SuperSignal West Pico PLUS Chemiluminescent Substrate (Thermo Fisher Scientific cat#: 34580) and captured with an ImageQuant LAS 4000 machine. To detect additional proteins, membranes were treated with Restore Western Blot Stripping Buffer (Thermo Fisher Scientific cat#: 21059), blocked, and then incubated with primary antibody. Antibodies against SCD (ab19862, 1:1000) and ATF4 (ab184909, 1:1000) were purchased from Abcam. Antibodies against PERK (3192, 1:1000), p-eIF2α (3597, 1:1000), eIF2α (9722, 1:1000), CHOP (2895, 1:1000), caspase-3 (14220, 1:1000), and cleaved caspase-3 (9664, 1:1000) were from Cell Signaling Technology. Antibody against GAPDH (H86504M, 1:20,000) was from Meridian Life Science, Inc. Antibody against β-actin (A1978, 1:20,000) was from MilliporeSigma. GAPDH or β-actin was used as loading control.

**Immunohistochemistry (IHC).** SCD protein was detected by IHC staining on ovarian cancer and fallopian tube paraffin-embedded sections of a tissue array purchased from the Cooperative Human Tissue Network (CHTN). The OVCA2 TMA contains 12 cases of serous papillary carcinoma, 12 cases of clear cell carcinoma, 12 cases of endometrioid adenocarcinoma and 12 cases of mucinous adenocarcinoma, 6 cases of serous borderline tumor and 6 cases of mucinous borderline tumor. Non-malignant tissue types were also included, namely 6 cases of proliferative endometrium, 6 cases of fallopian tube fimbriae, 6 cases of ovarian serous cystadenoma and 6 cases of ovarian mucinous cystadenoma. Each specimen is represented by 4 replicate cores on the array. The array slide was heated at 56°C for 30 minutes to melt the paraffin. Removal of paraffin was completed using xylene and then sections were rehydrated by immersing the array in decreasing concentrations of ethanol solutions. Heat-induced antigen retrieval was performed with 10mM sodium citrate buffer (pH = 6.0) at 95℃ for 30 min, followed by cool down at room temperature for 1 hour. Sections were treated with 3% hydrogen peroxide for 15 minutes, washed with PBS, blocked with 3% normal goat serum to decrease non-specific binding, and then incubated with anti-SCD antibody (5µg/mL; ab19862, Abcam), overnight at 4°C. Mouse IgG was used as negative control (sc-2025, Santa Cruz Biotechnology) on an ovarian cancer xenograft run in parallel. Sections were incubated with biotinylated secondary antibody for 15 minutes and then with streptavidin-HRP of a Dako (K0675, Agilent Technologies) and continued with incubation with Dako DAB+ Substrate Chromogen System (K3467, Agilent Technologies) to generate colored signal. Sections were counterstained with hematoxylin and imaged with a digital camera attached to a compound microscope. SCD staining was scored by a board-certified pathologist (JJW) recording the level of staining intensity (0, 1+, 2+, or 3+) for each tumor section. Each tumor or normal specimen was represented by 4 cores, the average of staining intensity for the 4 cores was calculated for the analysis.

**Lipidomics.** OVCAR-5 cells transduced with shRNA targeting SCD vs. control shRNA cells were seeded at a density of 400,000 cells on a 10cm dish, cultured in regular media for 48h, and then in medium containing low serum (1% FBS) for an additional 48 hours. Cells were washed twice with ice cold PBS, harvested in 800µL ice cold PBS and spun down at 1000 rpm at 4℃ for 5min. Supernatants were discarded and cell pellets were frozen on dry ice and stored at -80°C until used for analysis at the Bindley Bioscience Center, Purdue University. Lipidomic analysis was performed using multiple reaction monitoring (MRM) profiling which has been designed to provide rapid and comprehensive analysis of several classes of lipids in biological samples (3). The MRM Profiling method workflow includes direct injection of diluted lipid extracts and the selection of signature fragments of lipid classes in a triple quadrupole mass spectrometer. Briefly, lipids were extracted using the Bligh & Dyer (1959) method (4). Dried lipid extracts were diluted in the injection solvent (acetonitrile/methanol/ammonium acetate 300mM 3:6.65:0.35 [v/v/v]) to obtain a stock solution. The stock solution was further diluted into injection solvent spiked with 0.1ng/µL of EquiSPLASH LIPIDOMIX (Avanti Polar Lipids cat#: 330731) for sample injection. The MRM-profiling methods and instrumentation used have been recently described in previous reports (5-10). Data acquisition was performed using flow-injection (no chromatographic separation) from 8µL of the diluted lipid extract stock solution delivered using a micro-autosampler (G1377A) to the electrospray ionization (ESI) source of an Agilent 6410 triple quadrupole mass spectrometer (Agilent Technologies, Santa Clara, CA, USA). A capillary pump was connected to the autosampler and operated at a flow rate of 7µL/min and pressure of 100bar. Capillary voltage on the instrument was 5kV and the gas flow 5.1L/min at 300°C. The MS data obtained from these methods was processed using an in-house script to obtain a list of MRM transitions with their respective sum of absolute ion intensities over the acquisition time. For the reporting of the relative amounts using normalization by the internal standards, the amount of each fatty acid was expressed as pg/1000 cells. For lipidomics profiling, cells were lysed in ultrapure water for lipid extraction. Statistical analysis was performed utilizing MetaboAnalystR 3.0 (11). Data on the relative amounts from different lipid classes were scaled to obtain a normal distribution, and evaluated by univariate analysis, principal component analysis (PCA), and cluster analysis/heatmap. Informative lipids were analyzed according to class, fatty acyl residue chain unsaturation level (12).

**RNA sequencing (RNA-seq).** RNA was extracted using TRI reagent. Genomic DNA was removed from 1µg total RNA with an RNeasy MinElute Cleanup Kit (QIAGEN cat#: 74204). Thereafter, mRNA was isolated from total RNA using a NEBNext Poly(A) mRNA Magnetic Isolation Module (New England Biolabs cat#: E7490S). This was followed by first strand cDNA synthesis, second strand cDNA synthesis, adaptor ligation, PCR enrichment, and purification for RNA-seq library preparation (New England Biolabs cat#: E7770S). RNA-seq library quality was determined with a BioAnalyzer. RNA-seq libraries were sequenced on an Illumina NextSeq 500 machine producing single-end reads with 75bp read length. Illumina software called bcl2fastq (v2.17.1.14) was used to convert base call files (*.bcl) to FASTQ format (*.fastq) with the following parameters: -r 4 -d 3 -p 8 -w 4. Raw RNA-seq reads were aligned to the human reference genome GRCh38 ENSEMBL release 91 (13) using STAR (v2.5.2) (14) and SAMtools (v1.6) (15). Mapped reads were then counted using HTSeq (Anaconda v3.6) (16). Differentially expressed (DE) genes between experimental groups were determined and FDR corrected for multiple hypothesis testing with the edgeR package (17) in R. Pathway analysis based on the differentially expressed genes was performed using Ingenuity Pathway Analysis (IPA) software (QIAGEN). Normalized counts for all genes and all biological replicates were exported from R and subject to Gene Set Enrichment Analysis (18) and Gene Ontology analysis using clusterProfiler (19).

### Supplemental figures

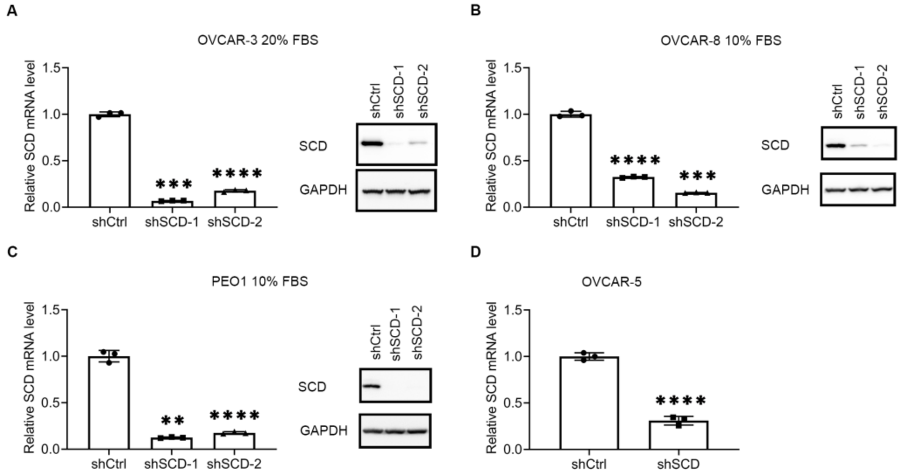


**Fig. S1.** (A-C) *SCD* expression measured by qRT-PCR (mean ± SD, n = 3) and western blot in OVCAR-3 (A), OVAR8 (B) and PEO1 (C) cells transduced with shRNAs (1 or 2) targeting *SCD* (shSCD), or control shRNA (shCtrl). (D) *SCD* expression measured by qRT-PCR (mean ± SD, n = 3) in shCtrl and shSCD OVCAR-5 cells cultured under low serum condition for 48 hours. * $p < 0.05$, ** $p < 0.01$, *** $p < 0.001$, **** $p < 0.0001$.

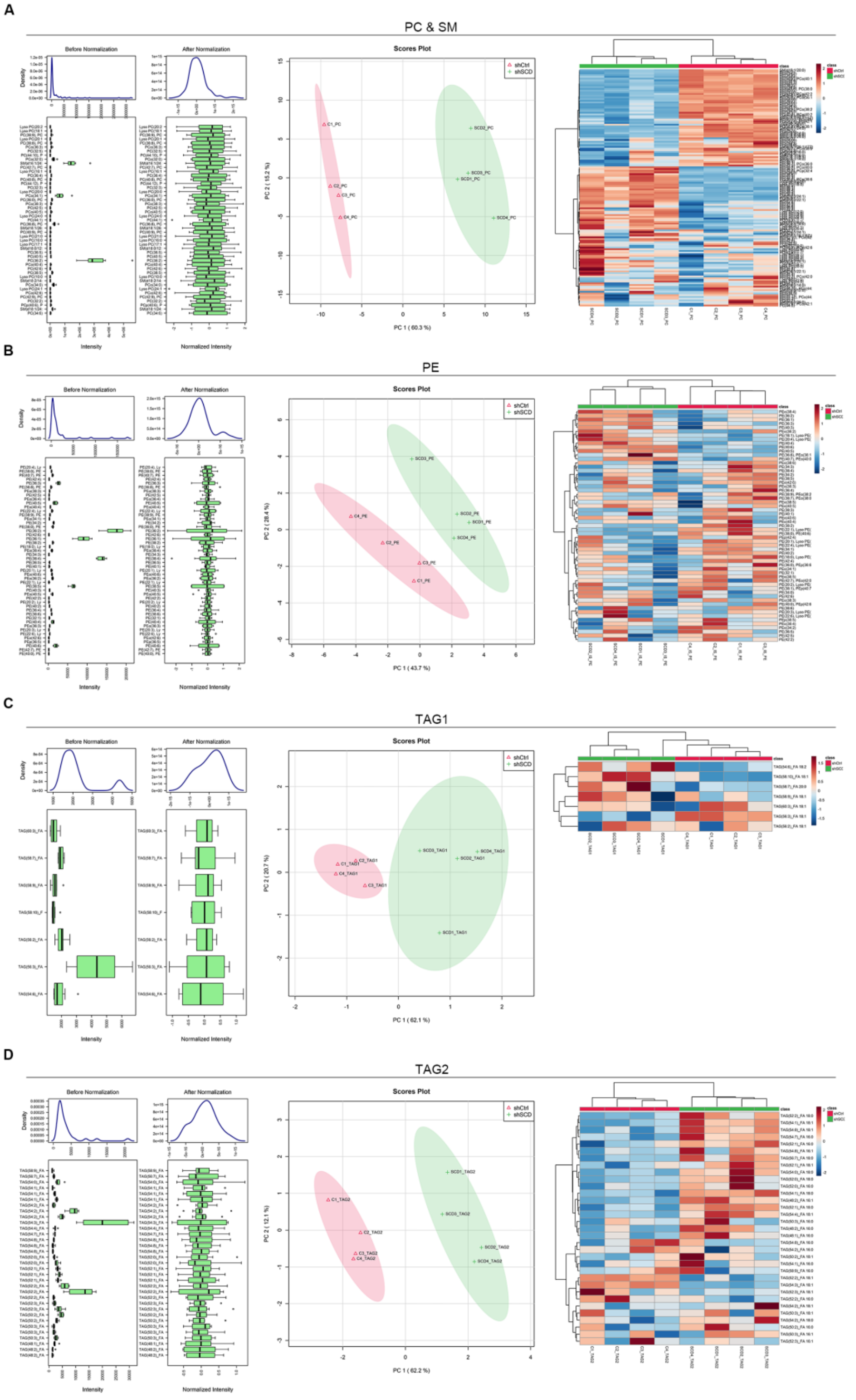
A
PC & SM
Before Normalization
After Normalization
Scores Plot
shCtrl
shSCD
PC 1 ( 60.3 %)
PC 2 ( 15.2 %)
B
PE
Before Normalization
After Normalization
Scores Plot
PC 1 ( 43.7 %)
PC 2 ( 28.4 %)
C
TAG1
Before Normalization
After Normalization
Scores Plot
PC 1 ( 62.1 %)
PC 2 ( 20.7 %)
D
TAG2
Before Normalization
After Normalization
Scores Plot
PC 1 ( 62.2 %)
PC 2 ( 12.1 %)

**Fig. S2.** (A-D) Normalization plot, principal component analysis and hierarchical clustering of phosphatidylcholine & sphingomyelin lipids (A), phosphatidylethanolamine (B), triacylglycerols 1 (C) and triacylglycerols 2 (D) from lipidomics profiling of shCtrl and shSCD OVCAR-5 cells cultured under low serum condition for 48 hours.

### General Statistics

Copy table | Configure Columns | Plot    Showing 6/10 rows and 3/9 columns.

| Sample Name | M Aligned | % GC | M Seqs |
|---|---|---|---|
| OVCAR5-shCtrl-1_S8_R1_001 | 49% | 32.6 | |
| OVCAR5-shCtrl-2_S13_R1_001 | 50% | 20.1 | |
| OVCAR5-shCtrl-3_S15_R1_001 | 50% | 29.3 | |
| OVCAR5-shSCD-1_S5_R1_001 | 49% | 28.2 | |
| OVCAR5-shSCD-2_S19_R1_001 | 51% | 18.3 | |
| OVCAR5-shSCD-3_S1_R1_001 | 51% | 29.3 | |

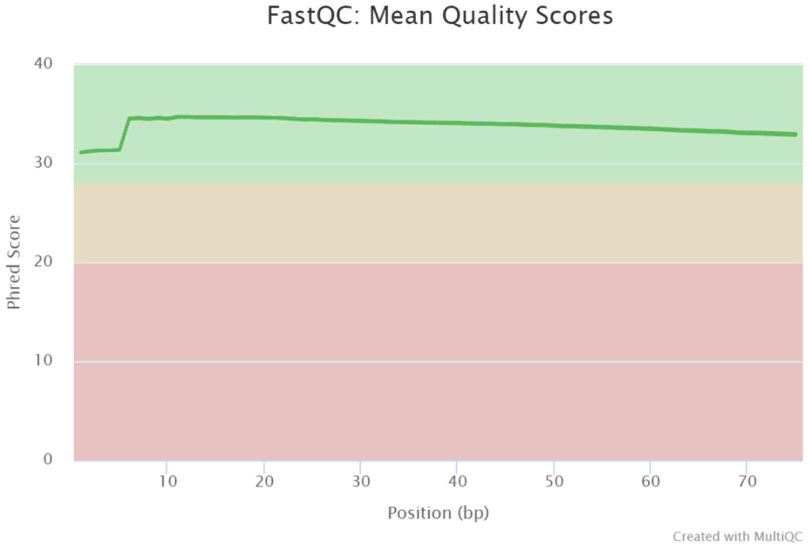


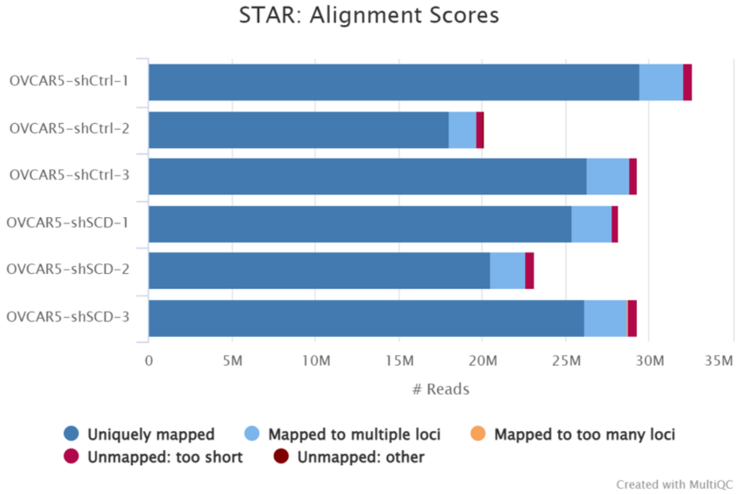


**Fig. S3.** Quality control of RNA-seq analysis of shCtrl and shSCD OVCAR-5 cells cultured under low serum condition for 48 hours.

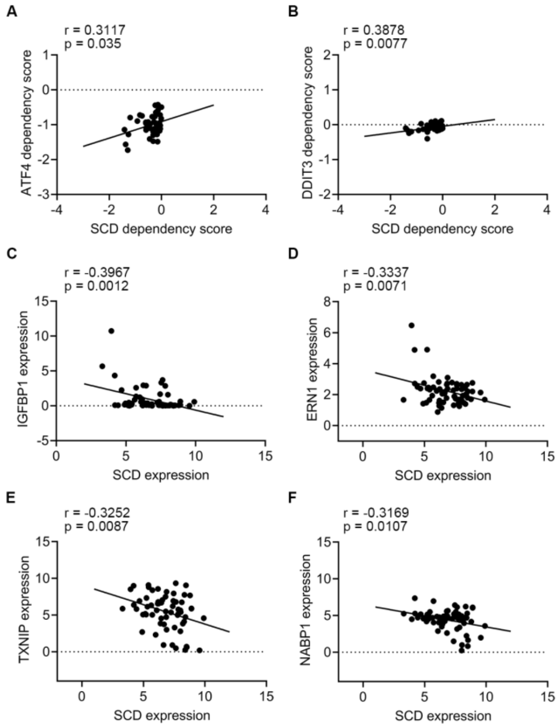


**Fig. S4.** (A, B) Pearson correlation analysis of dependency scores of SCD, ATF4 (A), and DDIT3 (B) in ovarian cancer cell lines (n = 46 cell lines). (C-F) Pearson correlation analysis between mRNA expression of SCD with IGFBP1 (C), ERN1 (D), TXNIP (E), and NABP1 (F) in ovarian cancer cell lines (n = 64 cell lines).

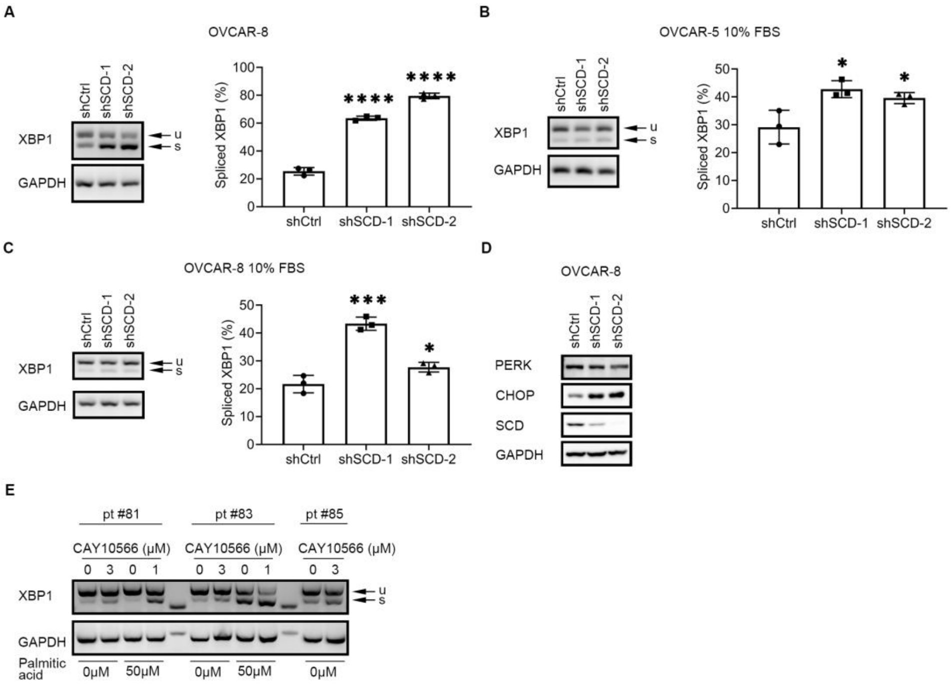


**Fig. S5.** (A) XBP1 splicing (u, unspliced transcript; s, spliced transcript) measured by RT-PCR and agarose-gel electrophoresis in OVCAR-8 cells transduced with control shRNA (shCtrl) or shRNAs (1 or 2) targeting SCD (shSCD) and cultured in low serum conditions (1% FBS) for 48 hours. Densitometric analysis of XBP1 splicing products is shown on the right. Bars represent percent of spliced XBP1 relative to total XBP1 (mean ± SD, n = 3). (B) XBP1 splicing (u, unspliced transcript; s, spliced transcript) measured by RT-PCR and agarose-gel electrophoresis in OVCAR-5 cells transduced with control shRNA (shCtrl) or shRNAs (1 or 2) targeting SCD (shSCD) and cultured in full serum conditions for 48 hours. Densitometric analysis of XBP1 splicing products is shown on the right. Bars represent percent of spliced XBP1 (mean ± SD, n = 3). (C) XBP1 splicing (u, unspliced transcript; s, spliced transcript) measured by RT-PCR and agarose-gel electrophoresis in OVCAR-8 cells transduced with control shRNA (shCtrl) or shRNAs (1 or 2) targeting SCD (shSCD) and cultured in full serum conditions for 48 hours. Densitometric analysis of XBP1 splicing products is shown on the right. Bars represent percent of spliced XBP1 (mean ± SD, n = 3). (D) Western blot of SCD and proteins of the PERK/eIF2α/ATF4 axis in OVCAR-8 shCtrl and shSCD cells cultured in low serum medium for 48 hours. (E) XBP1 splicing (u, unspliced transcript; s, spliced transcript) in primary cells from tumors of three ovarian cancer patients cultured in low serum conditions and treated with 3μM CAY10566 (SCD inhibitor) for 48 hours, or with 1μM CAY10566 and 50μM palmitic acid for 12 hours. * $p < 0.05$, ** $p < 0.01$, *** $p < 0.001$, **** $p < 0.0001$.

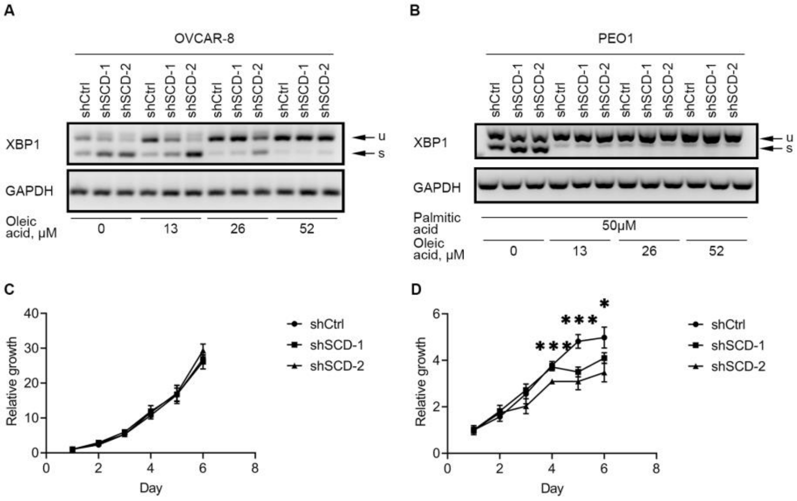


**Fig. S6.** (A) XBP1 splicing (u, unspliced transcript; s, spliced transcript) measured by RT-PCR and agarose gel electrophoresis in OVCAR-8 cells transduced with control shRNA (shCtrl) or shRNAs (1 or 2) targeting SCD (shSCD), cultured in medium containing low serum, and treated with indicated doses of oleic acid for 48 hours. (B) XBP1 splicing (u, unspliced transcript; s, spliced transcript) measured by RT-PCR and agarose gel electrophoresis in PEO1 cells transduced with control shRNA (shCtrl) or shRNAs (1 or 2) targeting SCD (shSCD), cultured in medium containing low serum, and treated with 50μM palmitic acid and indicated doses of oleic acid for 12 hours. (C) Proliferation curve of OVCAR-5 cells stably transduced with shRNA targeting SCD or control shRNA and cultured under full serum (10% FBS) conditions. (D) Proliferation curve of OVCAR-5 cells stably transduced with shRNA targeting SCD or control shRNA and cultured under low serum conditions (1% FBS). * $p < 0.05$, ** $p < 0.01$, *** $p < 0.001$, **** $p < 0.0001$.

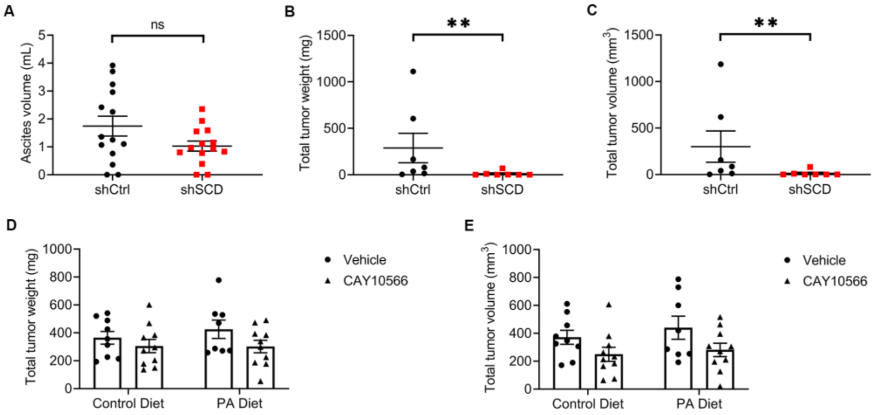


**Fig. S7.** (A) Ascites volume in athymic nude mice injected intraperitoneally with OVCAR-5 cells transduced with control shRNA (shCtrl) or shRNA targeting SCD (shSCD), and evaluated after 28 days (means ± SE, n = 14 per group). (B, C) Total tumor weight (B) and total tumor volume (C) in athymic nude mice injected subcutaneously with OVCAR-3 cells transduced with control shRNA (shCtrl) or shRNA targeting SCD (shSCD) (means ± SE, n = 7 per group). (D, E) Total tumor weight (D) and total tumor volume (CE) in athymic nude mice injected intraperitoneally with OVCAR-5 cells, fed with a palmitic acid-rich diet or control diet, and treated with SCD inhibitor CAY10566 or vehicle for 28 days. Values are means ± SE, n = 10. * $p < 0.05$, ** $p < 0.01$, *** $p < 0.001$, **** $p < 0.0001$.

**Table S1.** Summary of SCD immunohistochemistry staining in gynecologic tissue (graded from 0 to 3+).

| Tissue type | Total number of samples | Number of samples with score <= 1 | Number of samples with score >1 | P value# |
|---|---|---|---|---|
| Fallopian tube fimbriae | 6 | 6 | 0 | NA |
| Serous papillary carcinoma | 12 | 5 | 7 | 0.0377 |
| All ovarian cancers (serous, endometrioid, mucinous, clear cell) | 42 | 16 | 26 | 0.0061 |
| All non-malignant ovarian tumors (serous cystadenoma and borderline tumors) | 18 | 14 | 4 | 0.5392 |

#Statistical significance was calculated by using the Fisher's exact test comparing each group with the control (fallopian tube fimbriae)

**Table S2.** Information for the ovarian cancer tumors used in this study.

| Patient ID | Final Pathology Report | Tumor Origin |
|---|---|---|
| 81 | high grade serous carcinoma (stage IIIC) | omentum |
| 83 | high grade serous carcinoma (stage IIIC) | ovary |
| 85 | high grade serous carcinoma (stage IIIC) | omentum |

**Table S3.** Primers used for real-time RT-PCR.

| Target Gene | Forward (5'- 3') | Reverse (5'- 3') |
|---|---|---|
| *SCD* | ACGATATCTCTAGCTCCTATACC | GGCATCGTCTCCAACTTATC |
| *GAPDH* | GTATGACAACAGCCTCAAGAT | GTCCTTCCACGATACCAAAG |
| *XBP1* (splicing assay) | CCTGGTTGCTGAAGAGGAGG | CTCCAGAACTCCCCATGG |

**Dataset S1 (separate file).** OVCAR-5 SCD KD lipidomics profiling significant lipids species list.

**Dataset S2 (separate file).** OVCAR-5 SCD KD RNA-seq significant genes list.
